# MiRNA-223 Inhibits MouseGallstone Formation by Targeting Key Transporters in Hepatobiliary Cholesterol Secretion Pathway

**DOI:** 10.1101/2020.10.19.344580

**Authors:** Feng Zhao, Shiyu Ma, Wei Shen, Yanghao Li, Zihe Peng, Xiaolin Cui, Lina Xing, Xiang Li, Gang Liu, Lingling Jin, Tonghui Ma, Lei Shi

**Author notes:** Corresponding author. (L.Shi), or (TH. Ma). **Author contributions:** Research design: Lei Shi, Tonghui Ma Conducting experiments: Feng Zhao, Shiyu MA, Xiaolin Cui, Lina Xing, Zihe Peng, Xiang Li. Yanghao Li, Gang Liu Data acquisition: Feng Zhao, Shiyu Ma, lingling Jin Data analysis: Shiyu Ma, Feng Zhao Writing the manuscript: Lei Shi, Tonguhi Ma,Wei Shen.

## Abstract

**Backgroud:** MiRNA-223 has previously been reported to play an essential role in hepatic cholesterol homeostasis by suppressing cholesterol synthesis, attenuating cholesterol uptake by hepatocyte from and promoting cholesterol efflux into the blood. However, its role in regulation of biliary cholesterol secretion and gallstone formation remains unknown.

**Methods:** Mice with conventional knockout (KO), hepatocyte-specific knockout (ΔHepa) / knockdown (KD) or gain expression of miRNA-223 were included in the study and were subjected to lithogenic diet (LD) for various weeks. The gallbladders were harvested and subjected to cholesterol crystal imaging and gallstone mass measurement. Liver tissues were collected for western blotting, RT-qPCR, and immunohistochemistry staining. Levels of cholesterol, bile salt, phospholipids, and triglyceride were determined in serum, liver tissues, and bile by enzyme color reactive assays. 3’ UTR reporter gene assays were used to verify the direct target genes for miRNA-223.

**Results:** LD-induced gallstone formation was remarkably accelerated in miRNA-223 KO, ΔHepa, and KD mice with concurrent enhancement in total cholesterol levels in liver tissue and bile. Key biliary cholesterol transporters ABCG5 and ABCG8 were identified as direct targets of miRNA-223. Reversely, AAV-mediated hepatocyte-specific miRNA-223 overexpression prevented gallstone progression with reduced targets protein expression.

**Conclusion:** The present study demonstrates a novel role of miRNA-223 in the regulation of hepatic bile cholesterol secretion pathway and gallstone formation by targeting ABCG5 and ABCG8 expression. Therefore, elevating miRNA-223 would be a potentially novel approach to overcome the sternness of cholesterol gallstone disease.

**Graphical Abstract:** 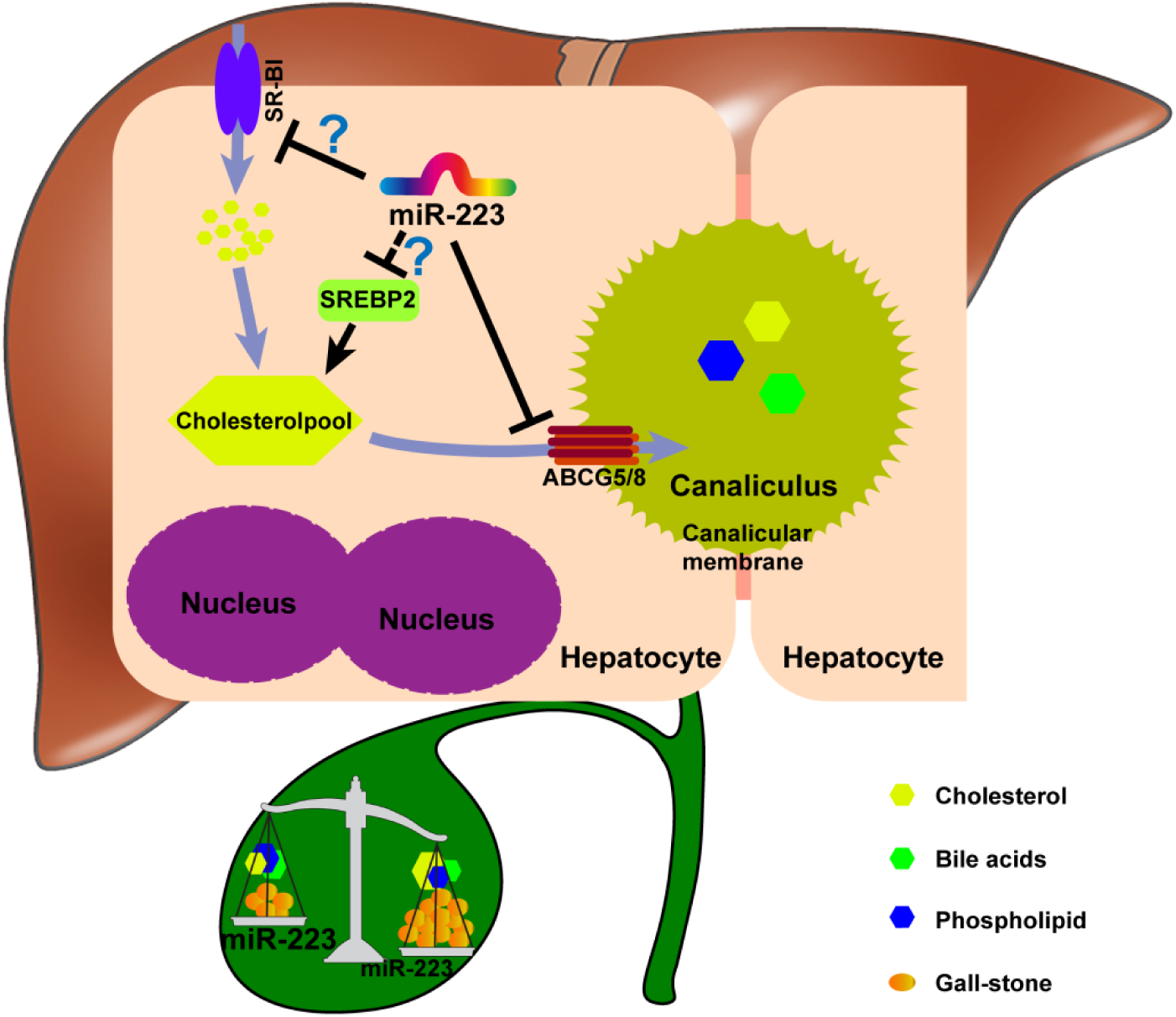

## Introduction

Gallstone is a disease of the digestive system with high incidence and approximately 10-20% prevalence in developed countries[1] with a gradual increase. The increasing annual cholecystectomy cases incur a huge economic stressand public health-care burden[2]. Based on chemical composition, cholesterol gallstone counts of 80-90% gallstone cases[2, 3] that appears due to imbalanced hepatic cholesterol metabolism associated with metabolic syndromes such as type 2 diabetes, obesity, hyperlipidemia, and high caloric intake. Dysregulation in genes controlling hepatobiliarycholesterol secretion has been shown to have a remarkable influence on cholesterol gallstone development. ABCG5 and ABCG8 are two heterodimeric ATP-binding cassette (ABC) transporters on hepatic canalicular membrane and are major contributors of hepatocytes cholesterol secretion into bile [4], as evidenced by approximately 75% reduced biliary cholesterol secretion rate in *Abcg5* and/or *Abcg8* KO mice [5]. A genome-wide association study (GWAS) revealed that genetic *ABCG5/8 loci* mutations are closely associated with gallstone diseases [1]. In *Abcg5/8* KO mice, LD-induced gallstone formation was greatly attenuated [6]. Other than ABCG5/8, scavenger receptor class B type I (SR-BI), coding by *Scarb1* in mice, has also been identified for efficient conduction of biliary cholesterol secretionin in physical condition [7]. Adenovirus-mediated hepatic overexpression of SR-BI has preferentially increased its canalicular membranes distribution and resulted in accelerated biliary cholesterol secretion rate in wild type (WT) or *Abcg5* KO mice [8]. In contrary, *Scarb1* KO or *Scarb1/Abcg 5* double KO mice exhibited reduced biliary cholesterol secretion [5]. Besides, SR-BI on hepatocyte basolateral membrane is well accepeted to conduct the cholesterol uptake from circulating HDL that plays a key role in maintaining serum cholesterol homeostasis [8, 9]. Therefore, these hepatic cholesterol transporters constitute an efficient cholesterol biliary secretion pathway and are proposed to be potential therapeutic targets for gallstone diseases.

The miRNAs play an important role in modulating pathophysiological processes in a fine-tuning manner by suppressing their target mRNA translation. In the liver, miRNAs are reported to actively contribute to metabolic homeostasis of glucose [10, 11], lipids as well as cholesterol [12, 13]. miRNA-223 gene is localized on the X chromosome and was initially identified as a hematopoietic specific miRNA that manipulates granulocyte differentiation [14], macrophage phenotype transition in the context of obesity [15, 16], and platelets ultra-activation under the diabetic situation [17]. As to specific liver pathological events, miRNA-223 was able to prevent APAP-induced liver damage via modulating neutrophil-mediated death-associated molecular pattern (DAMP) [18]. Moreover, miRNA-223 limits the neutrophil oxidative stress through suppressing IL-6–p47^phox^ pathway and protects the liver from alcoholic injury [19]. Notably, Vickers *et al* [13] reported that elevated intracellular cholesterol increases hepatocyte miRNA-223 expression and influences hepatic cholesterol homeostasis by different mechanisms i.e. attenuating hepatocyte cholesterol uptake from blood by targeting SR-BI expression, suppressing cholesterols synthesis via targeting HMGCS1 and SC4MOL, and promoting cholesterol efflux by indirectly upregulating basolateral ABCA1 expression in hepatocytes. Given that, we further wonder whether or not miRNA-223 also influences biliary cholesterol secretion pathway, the other dirction of liver processing cholesterol in addition to blood, and further consequences in gallstone formation.

The purpose of the present study is to determine the significance of hepatocyte miRNA-223 in gallstone pathogenesis with a special focus on its role in regulating the expression of key transporters in the biliary cholesterol secretion pathway.

## Methods

### Animals

MiRNA-223 KO and conditional KO mice (cKO) were generated in C57BL/6J line by Shanghai Biomodel Organism Science & Technology Development Co., Ltd. Conventional, hepatocyte-specific, and myeloid-specific miRNA-223 KO mice were achieved by crossbreeding miRNA-223 cKO with Alb-Cre^Tg^ and Lyz2-Cre^Tg^ mouse lines provided by Biomodel Organism Science & Technology Development Co., Ltd. Adeno-Associated Viruses (AAVs) used in this study were purchased from Hanbio Biotechnology Co., Ltd. For hepatocyte-specific knockdown of miRNA-223 expression, AAV8-TBG-Flag-Cre-T2A-GFP (1×10^11^ virus genome) or AAV8-TBG-GFP (as control, 1×10^11^ virus genome) was injected intravenously via tail vein into miRNA-223 cKO mice. For hepatocyte-specific overexpression of miRNA-223 precursor sequence, wild-type (WT) mice were received a one-time injection of AAV8-U6-miRNA-223 precursor/CMV-GFP or AAV8-CMV-GFP (each 1×10^11^ virus genome) via the tail vein.

### Murine Gallstone Model

The 8 weeks old male mice with C57BL/6J genetic background were used in the study and fed with lithogenic diet (TP2890, containing 15% lard fat, 1.25% cholesterol, and 0.5% sodium cholic acid) or chow (LAD0011) purchased from TROPHIC Animal Feed High Tech Co. Ltd, China for the indicated time periods.

### Bile Flow Rate Measurements

The procedure was as described in previous publication[20]. Brifely, mice were fasted for 6 h allowing free access to water. After ligating the cystic duct, the common bile duct was cannulated with PE-10, and hepatic bile was collected for 30 min to determine the flow rate. During surgery and hepatic bile collection, the mice were under anesthesia with isoflurane and maintained at 37°C on a hot plate.

### Study Approval

Animals were maintained in Dalian Medical University Laboratory Animal Center under the specific pathogen-free condition, and all animal study procedures were approved (#AEE17036) by the Ethics Committee for Biology and Medical Science of Dalian Medical University.

### Cell Culture

Mouse primary hepatocytes were isolated by gradient density centrifugation of 30% Percoll solution after liver perfusion using Collagenase II (Sigma). Primary hepatocytes were cultured in William’s E Medium (Thermo Fisher Scientific) supplemented with 10% fetal bovine serum, 1% ITS (Sigma), 2 mM L-glutamine, and 100 nM dexamethasone. HEK293T cells were maintained in Dulbecco’s Modified Eagle’s Medium supplemented with 10% fetal bovine serum plus penicillin (100 mg/mL) and streptomycin (100mg/mL). Cells were kept in a 5% CO_2_ and 95% air humidified incubator at 37°C.

For FACS analysis, after Collagenase II *in vivo* digestion, hepatic cell suspension or purified hepatocytes were incubated with indicted primary antibodies at room temperature for 20 min in the dark and different cell populations were sorted and further analyzed by MOFLO ASTRIOS^EQ^, BECKMAN COULTER.

### Plasmids and Antibodies

The partial 3’ UTR regions of mouse *Abcg5* (Genebank: NM_031884.2, Range 2095 to 2411) and *Abcg8* (Genebank: NM_026180.3, Range 2394 to 2772) were obtained by RT-PCR from liver tissue cDNA and subcloned into the pMIR-Reporter plasmid. The mutants for miRNA-223 binding sequences in 3’ UTRs of *Abcg5* and *Abcg8* were obtained by using the Fast Mutagenesis System (TRANSGEN BIOTECH). The primer sequences for cloning and mutation are all shown in Supplementary List 1. The antibodies usage and information are presented in Supplementary List 2.

### Biochemical Analysis

Total cholesterol (T-Chol. Catalog No. A111-1-1), high-density lipoprotein cholesterol (HDL-C Catalog no. A112-1-1), low-density lipoprotein cholesterol (LDL-C Catalog No. A113-1-1), total bile acid (TBA Catalog No. E003-2-1), and triglycerides (TG Catalog No. A110-1-1) concentrations were analyzed using Kits (Nanjing JIANCHENG Bioengenering, Najing, China) according to the manufacturer’s instructions. Phospholipid (PL) concentrations were quantified using Wako Kits (Catalog No. 296-63801, Osaka, Japan) according to the manufacturer’s instructions. The activities were measured using the Multi-Mode Microplate Reader (BioTek Synergy NEO). Bile cholesterol saturation index was calculated as described in the previous report [21].

### Bile Cholesterol Crystal Analysis and Gallstone Examinations

The contents of gallbladders were placed onto glass slides or centrifugal tubes after cholecystectomy, followed by measuring the weights of gallstone after removal of bile and dry by air. Bile cholesterol crystal was evaluated under polarized-light microscopy (Olympus BX63) and the crystal type was classified according to previous study [22, 23].Gallstone and KBr mixture was prepared at a ratio of 1:150 and the contents of cholesterol, bile acid, and phospholipid were further analyzed by Fourier Infra-redSpectrograph (Nicolet 6700 FlexFT-IR SPECTROMETER, Thermo Fisher Scientific).

### Histological Analysis

After scarification, animal livers were embedded in paraffin or optimal cutting temperature compound (OCT). Paraffin-embedded tissue sections were stained with hematoxylin and eosin (H&E) for morphological studies and immunohistochemical analysis using SR-BI antibodies. OCT-embedded tissue cryosections were incubated with rhodamine-phalloidin (Sigma) and DAPI for fluorescence analysis.

### Oil Red O Staining

Hepatic steatosis was assessed by Oil Red O staining in OCT-embedded tissue sections using the Oil Red O Stain Kit (Catalog No. G1260) from SolarbioLife Sciences according to the manufacturer’s protocol.

### Real-Time quantitative PCR and Genotyping

Total RNA was extracted from mouse liver tissues using the TRIzol (Invitrogen) according to the manufacturer’s instructions and cDNAs were reverse-transcribed using the FastKing RT Kit (TIANGEN). The cDNA samples were amplified by Real-time quantitative PCR (RT-qPCR) using SYBR Green qPCR Master Mix (Bimake) and *18S* was used as a standard reference. For genotyping, the genomic DNA from indicated tissues or primary cells were prepared by TIANamp Genomic DNA Kit (TIANGEN) and further identified by PCR using specific primers. The primer sequences used in this study are presented in Supplementary List 1.

### Western Blot

Mouse tissues or primary hepatocytes homogenates were subjected to SDS-PAGE, transferred to nitrocellulose (NC) membranes, and incubated with specific primary antibody. After washing, the membranes were incubated with anti-mouse Dylight 680 or anti-rabbit Dylight 800 secondary antibody (Abbkine). The membranes were then scanned using the Odyssey CLx Imaging System (LICOR), and the images were generated employing the Image Studio software.

### Luciferase Activity Assay

HEK293T cells (5 x 10^5^/well) were cultured in 24-well plates one day before transfection, and then co-transfected with 60 ng 3′-UTR luciferase reporter plasmids, 20 ng β-galactosidase plasmids, 20 nM miRNA-223 mimics or the negative controls by using Lipofectamine RNAi MAX Transfection Reagent (Thermo Fisher Scientific) according to the manufacturer’s instructions. Cells were harvested 24 h post-transfection and further subjected to luciferase and β-galactosidase activitydetection by using the D-Luciferin (BD Biosciences) and o-nitrophenyl-β-d-galactoside (ONPG), respectively. The firefly luciferase and β-galactosidase activities were measured using the Multi-Mode Microplate Reader (BioTek Synergy NEO). The results were normalized as the ratio of firefly luciferase activities to β-galactosidase activities.

### Statistical Analysis

Data are expressed as the mean ± S.E.M, and statistical evaluation was performed using Student’s t-test for unpaired data. The values of *p*<0.05 or less were considered statistically significant.

## Results

### Hepatocytes miRNA-223 Expression Is increased with LD-induced Gallstone Formation

To analyze the correlation of hepatic miRNA-223 expression with cholesterol gallstone formation, liver tissues and primary hepatocytes were analyzed from wildtype mice on chow or challenged with lithogenic diet (LD) for 5 weeks. Along with LD-induced gallstone generation (Figure 1A) miRNA-223 expression (miRNA-223-3p levels, as its other mature form miRNA-223-5p is undetectable in the livers) was significantly increased in both liver tissues and freshly isolated primary hepatocytes (HCs) (Figure 1B-C), although the absolute miRNA-223 expression level in primary hepatocytes only accounts for ∼0.07% of that in liver tissues (Figure 1C). Besides, a significant hepatic leukocytes infiltration in LD-treated livers was observed (as indicated by arrow shown in Figure 1D), which was further evidenced by increased hepatic Gr1/CD11b-positive cell population (Figure 1E-F) and mRNA expression of leukocytes markers *CD11b, Ly6g, Ly6c* and *Mpo* (Figure 1G). Therefore, it is assessed that hepatic nonparenchymal cells especially the infiltrating leukocytes may account for the dramatically higher miRNA-223 levels in normal and LD-treated livers as miRNA-223 was known to highly expressing in leukocytes[14].

**Figure 1.**
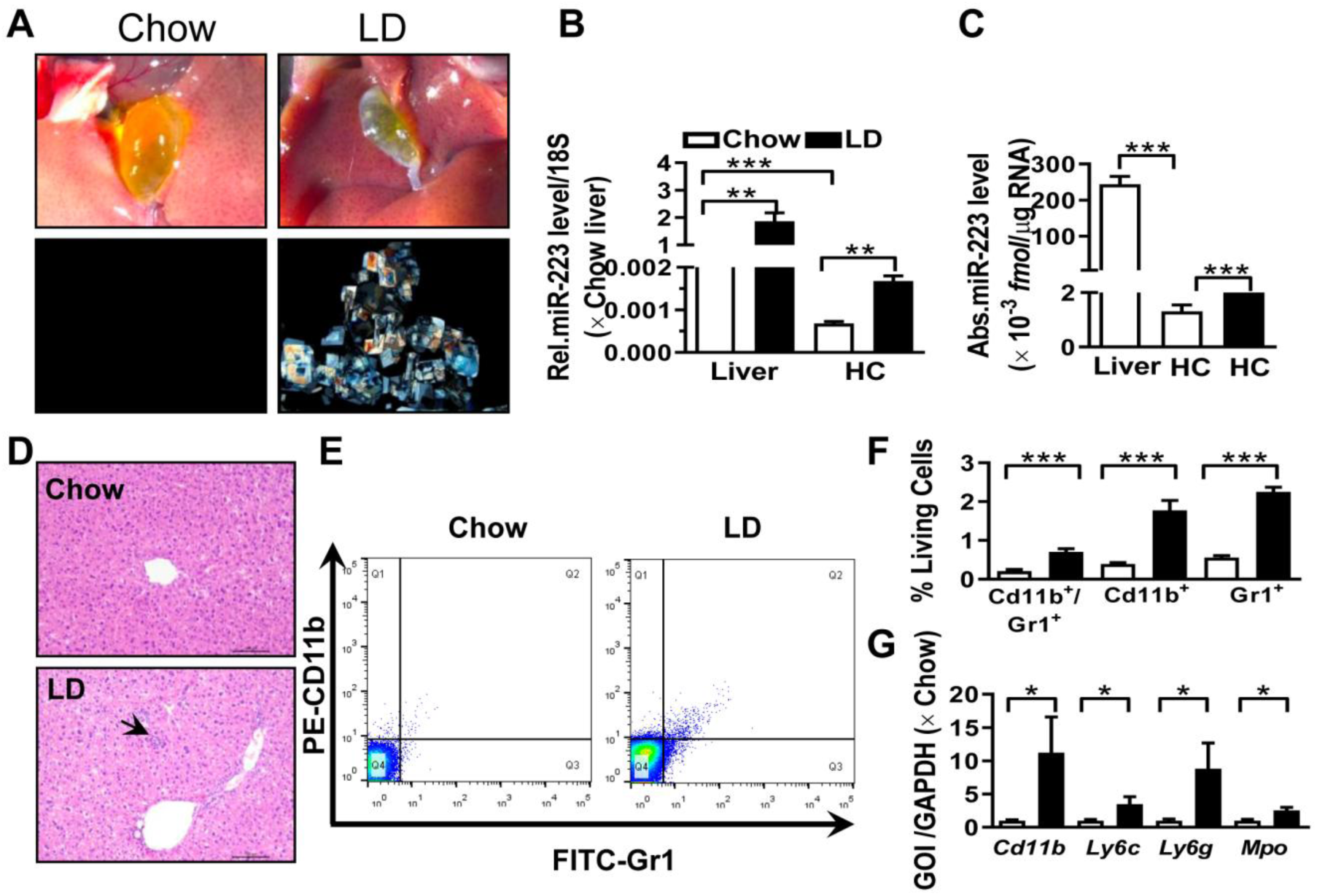
The expressional alternation of miRNA-223 in mouse liver during gallstone progression. WT male mice with 8-10 weeks were fed for lithogenic diet (LD) or chow diet for 5 weeks. A) Representative images showing the gall bladders and cholesterol crystals in bile; RT-qPCR detecting (B) the relative or (C) absolute miRNA-223 expression in liver tissues and primary hepatocytes (HC) from chow and LD fed mice with 18s as reference gene (n=5-6 mice per group); (D) H&E staining for liver sections; (E&F) FACS analysis for hepatic infiltrating CD11b^+^/Gr1^+^ cell population (n=3 mice per group); (G) Leucocytes marker genes were assessed by RT-qPCR (n=3 mice per group); *\*p*<0.05; \**\*p*<0.005; \*\**\*p*<0.001.

### MiRNA-223 Knockout Accelerates LD-induced Gallstone Formation in Mice

To determine the importance of miRNA-223 in gallstone formation, we firstly generated miRNA-223 conditional KO mice (miRNA-223 cKO, Supplementary Figure 1A) and further obtained conventional miRNA-223 KO mice (miRNA-223^-/y^ and -/y as a short form) via crossbreeding miRNA-223 cKO mice with EIIA-Cre^Tg^ mice (Supplementary Figure 1B). -/y was examined for the removal of miRNA-223 loci in liver genomic DNA (Figure 2A-B), which was further comfirmed by undetectable miRNA-223 expression in livers (Figure 2C). As shown in Figure 2D, -/y mice accelerated gallstone progression with more agglomerated cholesterol monohydrate crystals in the bile than that in WT littermates (miRNA-223^+/y^ and +/y as a short form). It was further evidenced by a greater gallstone forming rate one week post LD feeding (80% in -y *vs*. 20% in +/y, as given in Figure 2E) and increased gallstome mass after 5 weeks LD treatment (2.70 ± 0.85 mg in -/y *versus*. 0.90 ± 0.12 mg in +/y, as shown in Figure 2F). However, the mice of both genotypes represented a comparable growth curve as well as fat, muscle contents (Supplementary Figure2A-D).

**Figure2.**
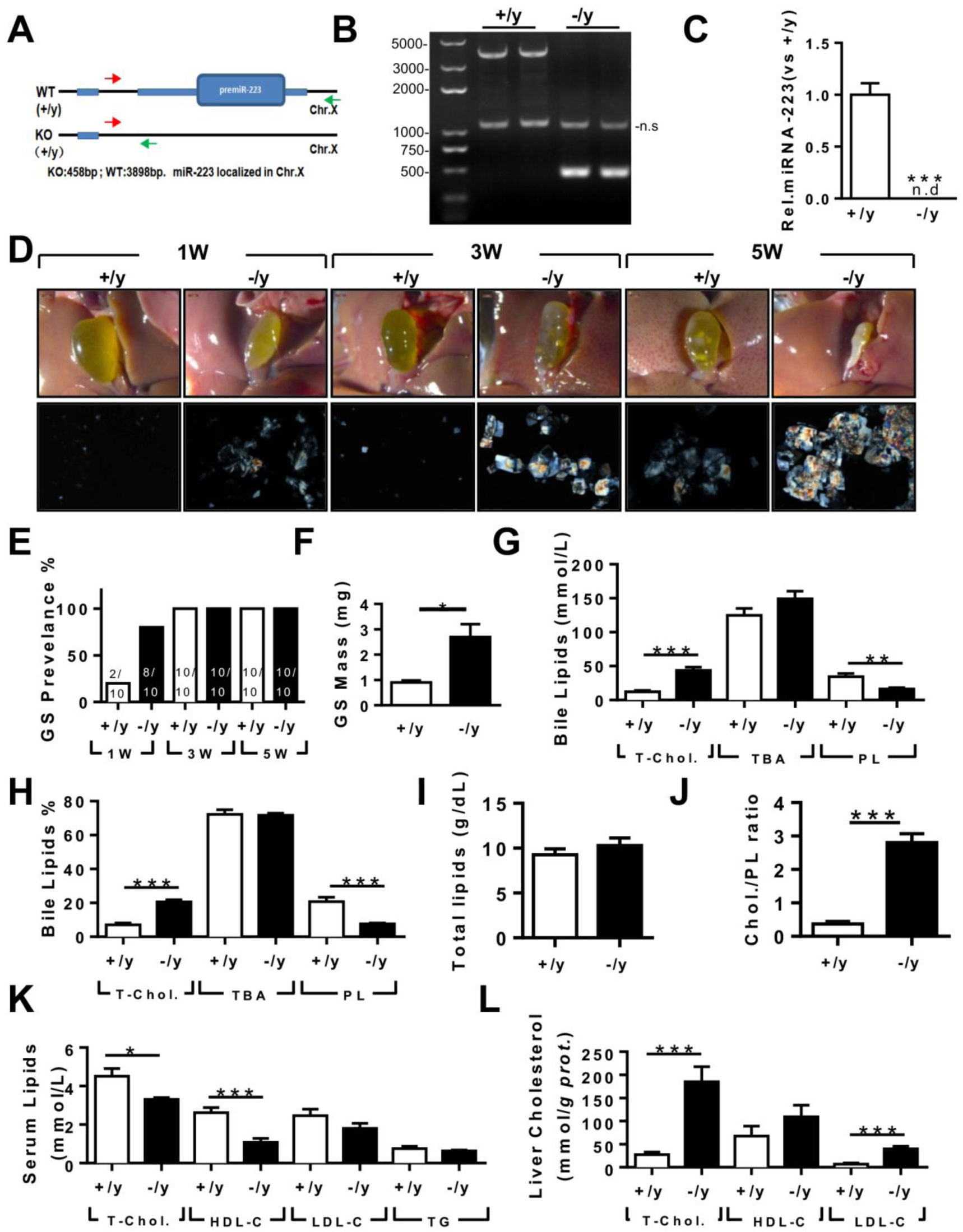
The effects of miRNA-223 KO on mice gallstone formation. MiRNA-223 KO (-/y) and WT (-/y) male littermates were subjected to LD feeding for 1, 3 and 5 weeks, and in the indicated time points, livers, serum and bile were harvested after overnight starvation for further detection. (A&B) Cartoon showing the miRNA-223 KO detection method by genomic DNA PCR, as miRNA-223 is localized in X chromosome, +/y and -/y were used to indicate miRNA-223 WT and KO respectively +/y: 3898bp, -/y:458bp, ns: non-specific bind; (C) RT-qPCR detecting miRNA-223 expression in livers, n=5 mice each group; (D) Representative images showing the gall bladders and cholesterol crystals in bile; (E) Gallstone prevalence with LD feeding summarized from 10 animals from each group; (F) weighted gallstone mass after 5 weeks LD treatment (n=5 mice per group); (G&H) bile lipids content and percentage; (I) total bile lipids contents; (J) ratio of cholesterol to PL in bile (n=6-8 mice per group); biochemistry parameters of Cholesterol, HDL-C,LDL-C and TG were separately determined in (K) serum and (L) liver tissues (n=10 mice per group). *\*p*<0.05; \**\*p*<0.005; \*\**\*p*<0.001versus +/y.

### MiRNA-223 Knockout Results in Cholesterol Super-Saturation in the Bile

In the bile, -/y mice exhibited significantly elevated levels of total cholesterol (T-Chol.) and reduced phospholipids (PL) content with no change in levels of total bile acid (TBA) (Figure 2G), resulting in a relative higher portion of cholesterol (20.67% in -/y vs 7.04% in +/y, as given in Figure 2H) and lower portion of PL (7.66% in -/y vs 20.70% in +/y, as presented in Figure 2H). Although there was no difference in total biliary lipids contents (Figure 2I), a remarkable elevationin of cholesterol / PL ratio in -/y mice were discovered (Figure 2J), indicating a super-saturation status for cholesterol. In parallel, significantly reduced serum T-Chol. and HDL-C (Figure 2K), enhanced liver T-Chol. and LDL-C levels (Figure 2L) and comparable liver TG contents (Supplementary Figure 2E) were also observed in -/y mice. There was no obvious liver injury as indicated by similar serum ALT and AST activities between the both –/y and +/y mice (Supplementary Figure 2F), but a slight increase in infiltrating inflammatory cells was observed in -/y livers (Suppymentary Figure 2G).

### Myeloid-specific miRNA-223 KO Has No Effect on Mouse Gallstone Formation

Myeloid miRNA-223 has been reported to profoundly influence cholesterol and TG metabolism in macrophages[13, 24] and hepatic immunoregulation [19, 25]. To rule out the possible involvement of myeloid miRNA-223 in cholesterol gallstone development, myeloid-specific miRNA-223 KO mice (ΔMye/y) were generated by crossbreeding the miRNA-223 cKO with Lyz-Cre^Tg^ mice (Supplementary Figure 1B). Precise genomic excision for miRNA-223 loci resulted in ∼98% reduction of miRNA-223 expression in ΔMye/y bone marrows (Supplementary Figure 3A&B). In ΔMye/y liver, we detected a dramatic decrease (∼78%) of miRNA-223 expression than that in cKO/y mice (Supplementary Figure3B), suggesting that the Lyz2-positive nonparenchymal cells contribute to a major portion of general hepatic miRNA-223 content. There was no obvious difference in major gallstone phenotypes (Supplementary Figure 3E-F) and liver injury (Supplementary Figure 3D), although a slightly lower levels of T-Chol. and LDL-C were detected in ΔMye/y serum (Supplementary Figure3C).

### Hepatocyte-specific Depletion of miRNA-223 Increases Gallstone Formation

To clarify the importance of hepatocytes miRNA-223 in LD-induced gallstone development, we generated hepatocyte-specific miRNA-223 KO mice (ΔHepa/y) by crossbreeding the cKO mice with Alb-Cre^Tg^ mice (Supplementary Figure1B). Interestingly, there was an approximately 97.35% decreased miRNA-223 expression was assessed in freshly isolated primary ΔHepa/y hepatocytes compared to that in cKO/y hepatocytes, whereras comparable levels of miRNA-223 were detected in livers tissues of both genotypes (Figure 3A). Upon five weeks LD treatment, ΔHepa/y remarkably promoted gallstone progression (Figure 3B). As early as one week after LD feeding, ΔHepa/y displayed a faster gallstone formation rate (90% in ΔHepa/y *vs*. 30% cKO/y, as given in Figure 3C), with a significant increase in gallstone mass (4.02±0.37 mg in ΔHepa/y *vs*. 2.4±0.38 mg in cKO/y, as presented in Figure 3D). Similarly, ΔHepa/y mice showed the alternations of lipids contents in bile (Figure 3E-H), serum (Figure 3I), liver tissue (Figure 3J, Supplementary Figure 4E) were greatly consistent with that appeared in -/y mice, except reduced biliary total lipids (Figure 3G) and comparable T-Chol. and elevated TG levels in serum (Figure 3J). Moreover, ΔHepa/y would not able to cause liver damage (Supplementary Figure 4A) when compared with cKO/y litermates, and it did not trigger further inflammatory cells infiltration, an observation discovered in -/y livers (Supplementary Figure 4C).

**Figure 3.**
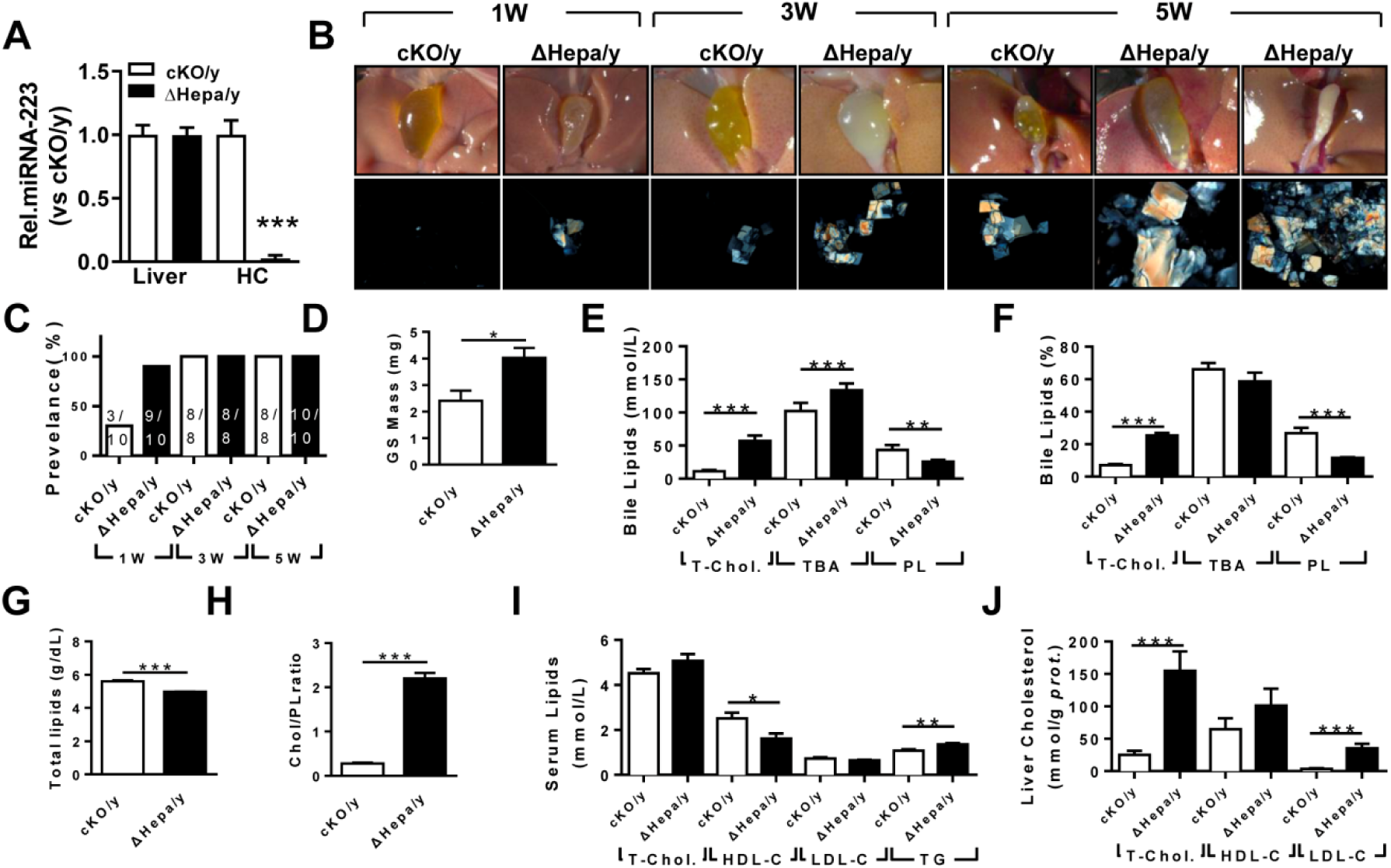
The influences of hepatocytes specific miRNA-223 KO on mice gallstone formation. Hepatic specific miRNA-223 KO (ΔHepa/y) and cKO/y littermates were subjected to LD feeding for 1 to 5 weeks, and in the indicated time points, livers, serum and bile were harvested after overnight starvation for further detection. (A) RT-qPCR showing miRNA-223 expression in livers or primary hepatocytes (n=5 mice each group); (B) Representative images showing the gall bladders and cholesterol crystals in bile; (C) Gallstone prevalence with LD feeding (n=8-10 mice each group); (D) Weighted gallstone mass (n=4 mice each group); (E&F) Bile lipids content and percentage; (G) total bile lipids contents; (H) ratio of cholesterol to PL in bile (n=8-10 mice each group); biochemistry parameters of Cholesterol, HDL-C,LDL-C and TG were separately determined in (I) serum and (J) liver tissues (n=8-10 mice each group); *\*p*<0.05; \**\*p*<0.005; \*\**\*p*<0.001versus cKO/y.

### AAV-Cre-mediated Knockdown of miRNA-223 in Liver Accelerates Gallstone Formation

To exclude potential compensatory effects from genetic depletion strategy, adenovirus associated virus (AAV)-mediated hepatocytes-specific knockdown (KD) of miRNA-223 expression (miRNA-223^KDHep^, KD_Hepa_) was performed by using Flag-tagged Cre recombinase under control of TBG promoter. Three weeks after infection, LD was applied for another 5 weeks (Figure 4A). The expression efficiency of Cre recombinase was visualized by immunofluorescent staining against Flag tag fusion expression with Cre (Figure 4B) and further verified by western blotting (Figure 4C). Consequently, liver genomic DNA excision for miRNA-223 loci (Figure 4D) caused an approximately 79.26% reduced miRNA-223 expression in freshly isolated hepatocytes (Figure 4E), although the general miRNA-223 amount in livers was not changed (Figure 4E). Consistently, KD_Hepa_ mice displayed similar phenotypes in gallstone promoting effects and lipids content changes as seen in ΔHepa/y mice (Figure 3F-K). Moreover, after KD_Hepa_, serum HDL-C levels were slightly reduced (Figure 4L) whereas liver T-Chol., LDL-C and TG were significantly enhanced (Figure 4M, Supplementary Figure 4F). Similarly, KD_Hepa_ also did not cause further liver injury and leukocytes infiltration (Supplementary Figure 4B&D).

**Figure 4.**
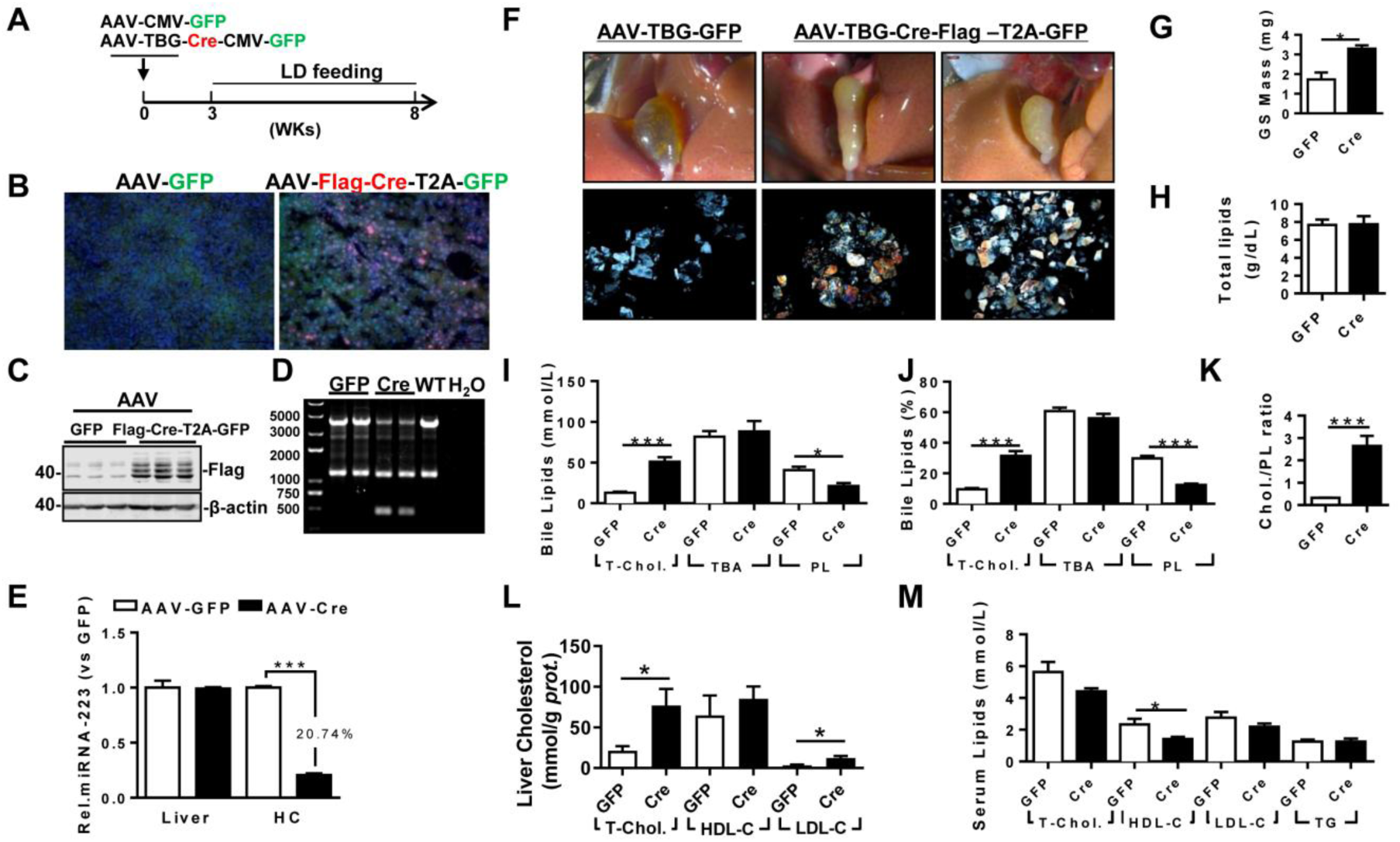
The influences of hepatocytes specific miRNA-223 KD on mice gallstone formation. (A) Hepatic specific miRNA-223 KD were conducted by injection miRNA-223 cKO micewith AAV-TBG-Flag-Cre or AAV-TBG-GFP for once(1×10^11^ virus genome) and 3 weeks later were feed with LD for additional 5 weeks, thereafter serum, livers and bile were harvested for further analysis. (B) Hepatic Flag-Cre overexpression were visualized by immunofluorescence against Flag tag in frozen liver sections (Flag, red; GFP, Green; DAPI, blue) and by (C) western blotting (∼42 kDa for Flag-Cre); The miRNA-223 KD efficacy was further determined by (D) hepatic genome excise (cKO.+/y:4130bp, KD: 458bp); (E) miRNA-223 expression in livers and HCs determined by RT-qPCR (n=4-6 mice per group); (F) Representative images showing the gall bladders and cholesterol crystals in bile; (G) weighted gallstone mass; (H) total bile lipids contents; (I&J) bile lipids content and percentage; (K) ratio of cholesterol to PL in bile (n=6-8 mice per group); biochemistry parameters of Cholesterol, HDL-C, LDL-C or TG were separately determined in (L) serum and (M) liver tissues (n=6-8 mice per group). *\*p*<0.05; \**\*p*<0.005; \*\**\*p*<0.001versus GFP.

Next, we examined the role of miRNA-223 deficiency in hepatocytes on bile secretion via a bile duct catheration method. As indicated in Supplementary Figure 5A&B, miRNA-223 KD_Hepa_ was not able to affect bile secretion speed both in chow or LD feeding conditions. However, total cholesterol level was significantly elevated from bile secreted from KD_Hepa_ livers with LD treatment (Supplementary Figure 5C), but levels of PL and TBA was comparable with AAV-GFP treated mice (Supplementary Figure 5D&E). Furethermore, Fourier Infra-red Spectrograph was used to analyze contents of cholesterol according to the transmission rate in specific absorption wavelength. As shown in Supplementary Figure 5F, relative lower transmission rate for cholesterol was observed in gallstone from AAV-Cre treatment mice, demonstrating higher cholesterol contents in the gallstone.

### MiRNA-223 Targets Key Transporters of Biliary Cholesterol Secretion Pathway

Based on the high concentration of biliary cholesterol secreted from hepatic miRNA-223 deficient mice, we postulated that miRNA-223 might expressionally or functionally affect key cholesterol transporters on the canalicular membranes. We detected the both protein and mRNA expression levels of ABCG5 and ABCG8, the major direct transporters for billary cholesterol secretion, were all significantly increased in ΔHepa/y livers in compare with that in cKO/y livers (Figure 5A-C). Besides, imperfect potential binding sequences of miRNA-223 on the 3’untranslational region (UTR) of murine *Abcg5* and *Abcg8* were discovered (Figure 5D). By using 3’UTR reporter gene assays, overexpression of miRNA-223 mimics was readily suppressing their WT reporter activities and the effects were eliminated when the seeding sequences were mutated (Figure 5E).

**Figure 5.**
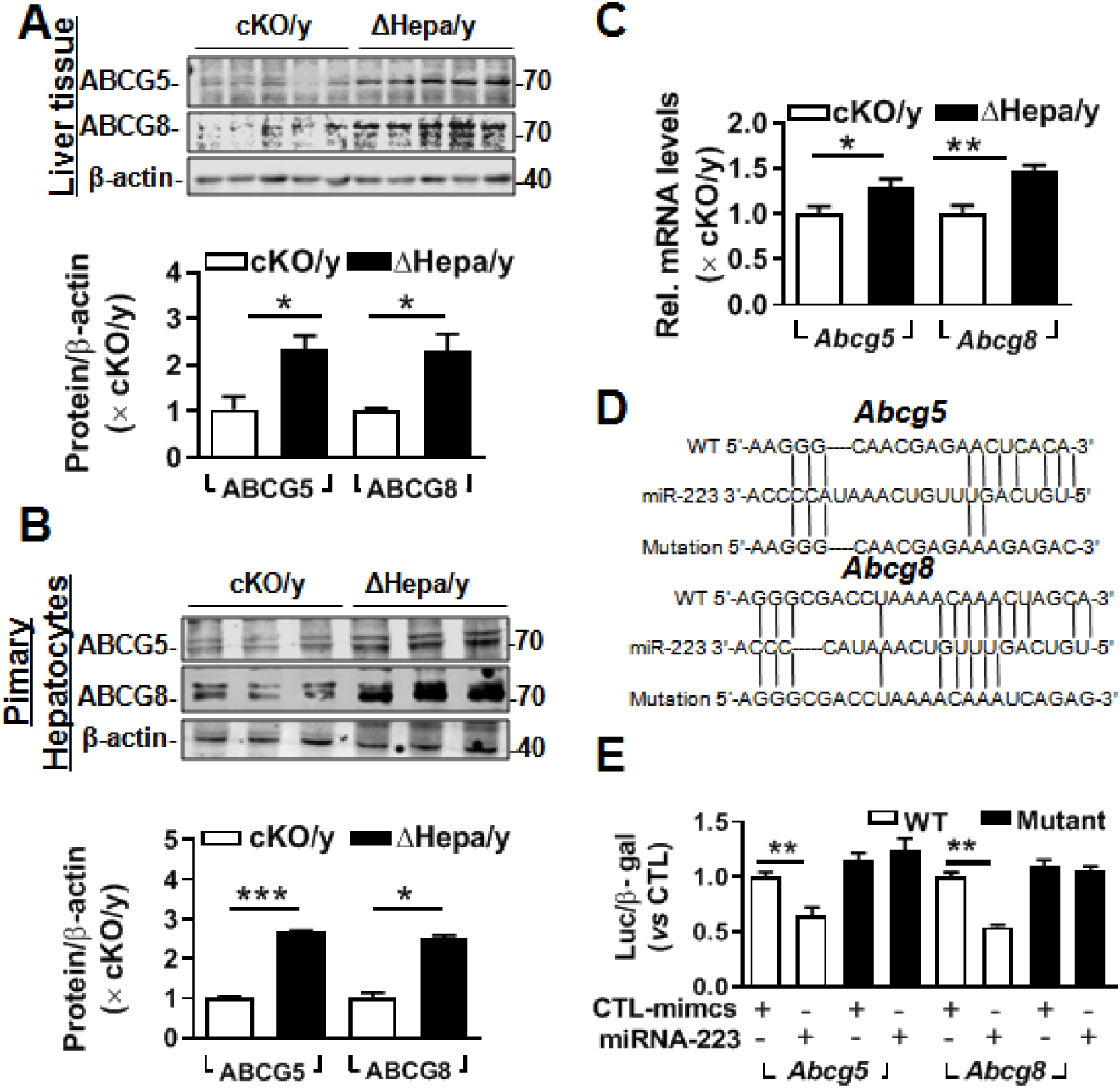
MiRNA-223 targets *Abcg5* and *Abcg8* expression in murine hepatocytes. After LD feeding for 5 weeks, ABCG5 and ABCG8 expression in ΔHepa/y and cKO/y livers were analysed in (A) protein levels by Western blotting and (C) mRNA by RT-qPCR (n=5 mice each group); (B) Protein levels of ABCG5 and ABCG8 from freshly isolated primary hepatocytes were examined by Westen blotting (n=3 mice per group). (D) Predicted binding sequences of miRNA-223 in 3’UTR regions of mouse *Abcg5* and *Abcg8* mRNA and (E) the effects of miRNA-223 mimics overexpression on luciferase activities determined from 3’UTR reporter assays for WT or muated forms of mouse *Abcg5, Abcg8* and *Scarb*1 (n=3 time indepemdent expreiments). *\*p*<0.05; \**\*p*<0.005; \*\**\*p*<0.001 versus cKO/y or control.

To further verify other potential mechanisms underlying the miRNA-223 deficiency-induced abnormalities in hepatic/biliary cholesterol homeostasis as a series of functional related genes were reported to be dysregulated in miRNA-223 KO livers[13]. We wondered how consistance of the effects of hepatocyte specific KO with conventional KO for miRNA-223 on those key genes expression in governing liver cholesterol or lipids metabolsim processes. Therefore, liver samples from -/y and +/y mice or ΔHepa and cKO/y littermates with 5 weeks LD treatment were further analyzed for gene expression in mRNA or protein levels. Hepatic cholesterol homeostasis is tightly harmonized by uptake, efflux, synthesis, and catabolism. For mRNA expression of cholesterol synthesis related genes, we found that *Srebp2* presented a consistent enhancement in -/y and ΔHepa/y livers, increases of *Hmgcs* and *Acat2* were only detected in ΔHepa/y livers, but *Sc4mcl* expression was not affected. For cholesterol uptake and efflux associated genes, *Abca1* mRNA was increased in both -/y and ΔHepa/y livers. *Sp3* mRNA but not *Sp1* was increased only from -/y livers. Similarly, increased *Vldlr* and decreased *Ldlr* mRNA levels were detected in -/y livers but not in ΔHepa/y livers. Besids, *Scarb1* mRNA expression were not able to link with miRNA-223 expression (Supplymentary Figure 6A). For hepatic bile acid synthesis related genes, a remarkably reduced *Cyp7a1* and *Cyp8b1* mRNA levels were detected in ΔHepa/y livers. In addition, we discovered an enhanced expression of *Abcb4* and *Abcb11* in ΔHepa/y livers. However, the mRNA levels of key transcriptional factors such as *Fxr, Rxra* were not impacted except slightly elevated *Lxr* (Supplymentary Figure 6E).

**Figure 6.**
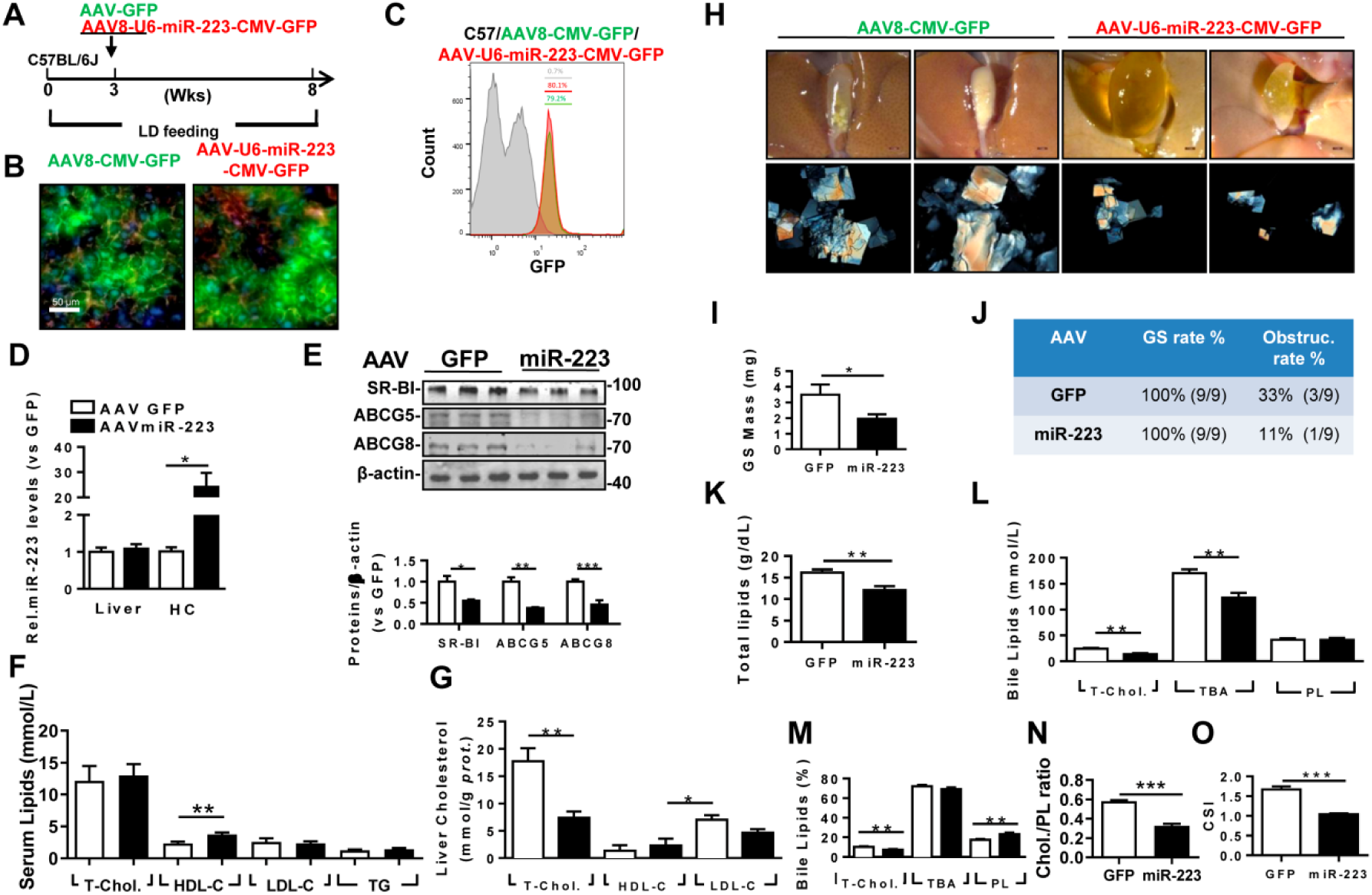
The therapeutic attempt against mice gallstone progression by specific OE of miRNA-223 in hepatocytes. WT mice were pretreated with LD for 3 weeks followed by (A) one-time injection with AAV8-U6-miRNA-223/CMV-GFP or AAV8-CMV-GFP (1×10^11^ virus genome) and continued LD feeding for additional 5 weeks, thereafter serum, livers and bile were harvested in the indicated time points for further analysis. AAV infection efficiency was assessed by (B) IF staining from frozen liver sections (GFP, green; DAPI, blue; F-actin, red) as well as by (C) FACS for GFP positive cells population from freshly isolated primary hepatocytes; (D) RT-qPCR evaluated miRNA-223 OE efficiency in liver and primary hepatocytes (n=3-5 mice per group); (E) the protein expression were examined in lives by Western blotting (n=3 mice per group); biochemistry parameters of total cholesterol, HDL-C, LDL-C and TG were determined in (F) serum and (G) liver tissues (n=9 mice per group); (H) Representative images showing the gall bladders and cholesterol crystals in bile; (I) Weighted gallstone mass (n=9 mice per group); (J) Table summarized gallstone formation rate and obstruction rate; (K) total bile lipids contents; (L&M) bile lipids contents and percentage; (N) ratio of cholesterol to PL as well as (O) cholesterol saturation index (CSI) (n=6-8 mice per group). (P)The cartoon summarized the role of miRNA-223 in balancing cholesterol homeostasis in hepatocytes and bile. *\*p*<0.05; \**\*p*<0.005; \*\**\*p*<0.001versus GFP.

As previous research indicated, miRNA-223 KO would change a series of mRNA expressions governing Lipid metabolism (GWAS Lipids traits) and cholesterol synthesis related genes (SREBPs targets) in mouse livers[13]. Therefore, we further examined these mRNA expressions in -/y and ΔHepa livers. As a consequence of increased expression of *Srebp2* in mRNA and protein levels (Supplymentary Figure 6A,C-D), the mRNA expression of major SREBPs responding genes was mildly augmented in ΔHepa/y livers. However, mRNA expressions for GWAS lipid traits related gene were largely unaffected (Supplymentary Figure 6B). Besids, we found the protein expression of ABCA1 and SR-BI were significantly elevated in ΔHepa/y livers (Supplymentary Figure 6 C&D).

### MiRNA-223 Overexpression in Hepatocytes Prevents LD-induced Gallstone Progression

To evaluate the therapeutic potential against gallstone progression by specificlyincreasing miRNA-223 levels in hepatocytes, administration of AAV8-U6-miRNA-223/CMV-GFP (OE), or AAV8-CMV-GFP (GFP) towards WT mice was conducted in the third week during an eight-weeks LD feeding frame (Figure 6A). The AAV infection rates were comparable as determined by GFP signals from frozen liver sections (Figure 6B) and ∼80% GFP positive primary hepatocytes in both groups by FACS analysis (Figure 6C). Approximately 20-fold increased miRNA-223 levels was detected in primary hepatocytes isolated from miRNA-223 OE mice than that in GFP mice while its general expression level in liver tissues was unaffected (Figure 6D). MiRNA-223 OE attenuated protein expression of ABCG5, and ABCG8 (Figure 6E), though their mRNA was not intensively affected (Supplymentary Figure 7A). Moreover, protein expression of SR-BI, SREBP2 and ABCA1 were all declined in primary hepatocytes isolated from miRNA-223 OE livers (Supplymentary Figure 7B). Further analysis indicated miRNA-223 OE resulted in comparable serum lipids levels with exception of slight increased HDL-C levels (Figure 6F). Meanwhile, remarkable reducd contents of T-Chol. LDL-C and TG were detected in miRNA-223 OE liver tissues (Figure 6G, Supplymentary Figure 7E). Importantly, miRNA-223 OE partially prevented further gallstone progression as evidenced by decreased appearance of the gallbladder stones (Figure 6H). In line with lower gallbladder obstruction rate (Figure 6J) in the miRNA-223 OE group (11.1%) compared to that in GFP group (33.3%), gallstone mass was also greatly attenuated (Figure 6I). Moreover, in the bile, miRNA-223 OE reduced total biliary lipids content (Figure 6K) as well as T-Chol. and TBA levels (Figure 6I&L). However, bile lipids composition analysis revealed relative lower percentage of cholesterol and higher percentage of PL in miRNA-223 OE mice (Figure 6M), leading to a declined cholesterol/PL ratio (Figure 6N) and attenuated cholesterol saturation index (CSI, shown in Figure 6O). Besides, AAV-miRNA-223 OE would not induce further liver damage as indicated by the similar ALT and AST activities in serum and normal liver structures (Supplementary Figure 7C&D).

**Figure 7.**
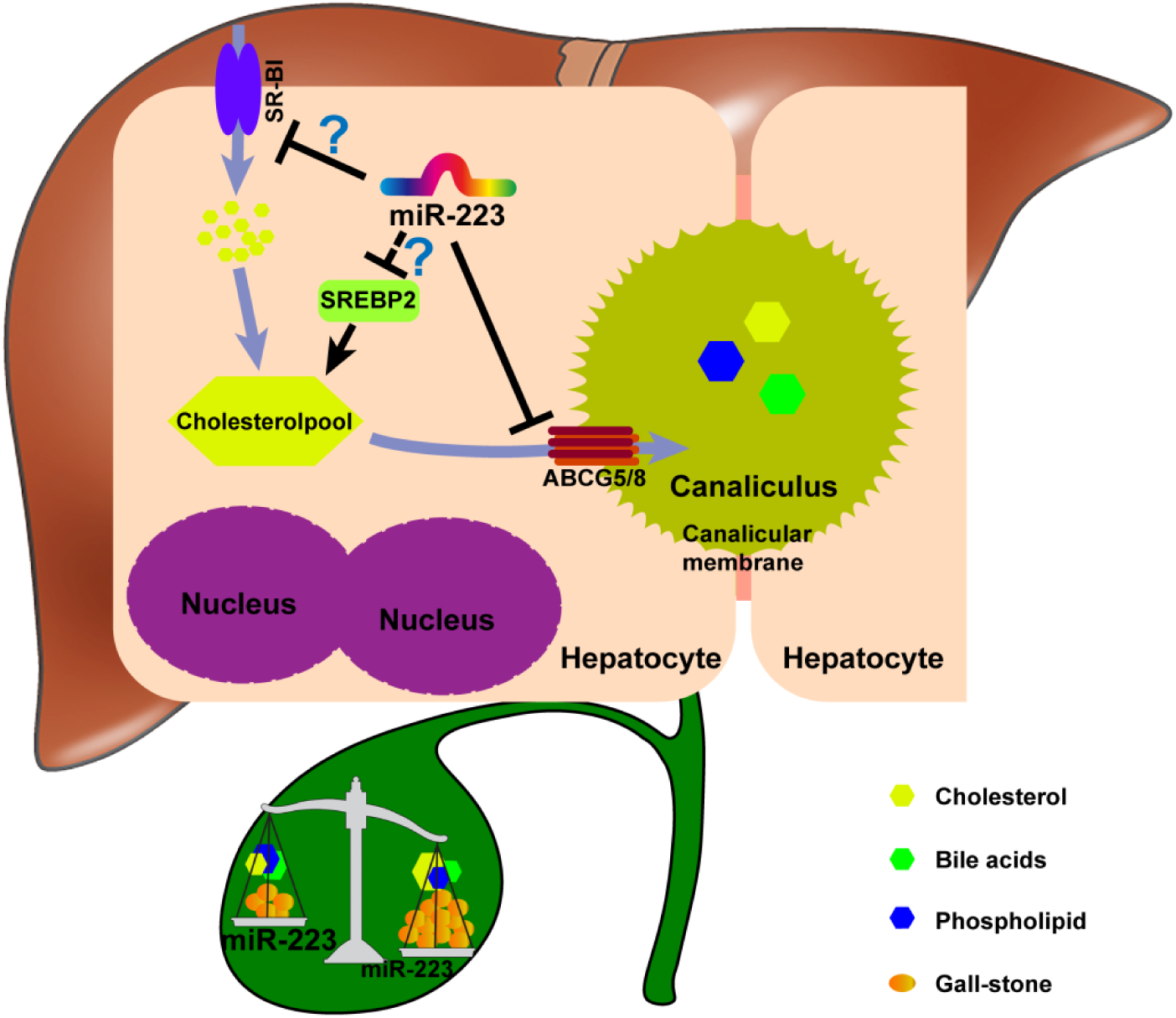
The potent role of miRNA-223 in regulating mice gallstone development.

Taken together, we proposed a novel mechanistic role of miRNA-223 in regulating LD-induced cholesterol gallstone development (Figure 7): miRNA-223 preferentially decreases cholesterol transportation from hepatocytes into the bile by directly targeting ABCG5 and ABCG8 at the canalicular membrane. The blockage effects of miRNA-223 on cholesterol biliary secretion pathway give rise to attenuate bile cholesterol content and saturation status, consequently prevent cholesterol gallstone formation. Besides, miRNA-223 may decline hepatic cholesterol synthesis by suppressing SREBP2 expression via an unclear mechanism. Moreover, by an identfyed manner, miRNA-223 would suppress SR-BI protein expression at basolateral membranes would inhibit hepatic cholesterol uptake (perheps decreaseing SR-BI expresion at canalicular membranes might precent biliary cholesterol secretion, a function confirmed in physical condtion but has not been verified in cholesterol gallstone pathogenesis yet.)

## Discussion

The purpose of this study was to determine the importance of hepatocytes miRNA-223 in regulating cholesterol biliary secretion and gallstone formation by using a series of genetically modified miRNA-223 transgenic mouse models. The major finding is that miRNA-223 plays a pivotal role in regulating hepatobiliary cholesterol secretion and cholesterol gallstone formation by targeting key cholesterol transporters.

We found that the LD with 1.25% cholesterol readily increases miRNA-223 expression in mouse liver tissue and freshly isolated primary hepatocytes. However, the increased miRNA-223 expression in liver tissue would be mainly accounted by infiltrated myeloid cells or Lyz-2 positive nonparenchymal cells because of ∼ 78% reduction of miRNA-223 expression was identified in liver tissue from myeloid specific miRNA-223 KO mice. Our data was suggestive for determing the importance of hepatic cell types in miRNA-223 mediated functional studies, especially for hepatocytes in which miRNA-223 expressional alternation would be carefully detected not only in liver tissues but also primary hepatocytes. Of course, we still can not exclude that LD may also influence miRNA-223 expression in other cell types in liver. A recent study reported that a high fat diet increased miRNA-223 expression in mouse liver and hepatocytes that consequently induced hepatic inflammation and fibrosis[26]. Cholesterol would be a key element to trigger miRNA-223 levels in hepatocyte as previous study implied cholesterol deprivation time dependently reduced its expression in human hepatocytes[13]. Moreover, we observed mild hepatitis and fibrosis in miRNA-223 KO liver treated with LD for 5 weeks. However, such abnormalities were not seen in hepatocyte-specific miRNA-223 knockout mice, suggesting that hepatocytes miRNA-223 is not involved in the development of pathological changes at least in current experimental settings.

To better determine the role of liver miRNA-223 in cholesterol gallstone formation, we generated a conditional miRNA-223 KO mice model and obtained conventional, hepatocytes-specific, and myeloid-specific KO mice by crossbreeding with EIIA-Cre, Alb-Cre, and Lyz-Cre transgenic mice. We also performed AAV-TBG-Cre-mediated hepatocytes-specific knockdown (KD_Hep_) in miRNA-223 cKO/y mice. By phenotypic analysis, we identified faster gallstone development in miRNA-223 conventional KO, ΔHep and KD_Hep_ mice but not in ΔMye mice when the animals were subjected to LD treatment, fully supporting the importance of hepatocytes miRNA-223 in regulating hepatic/biliary cholesterol homestasis. In a previous study using conventional miRNA-223 KO mice, Vickers *et al*. [13] found that miRNA-223 controls hepatic cholesterol uptake, biological synthesis, and efflux by targeting a complex gene set including *ScarbI, Hmgcs1, Abca1 etc*. In addition, they reported that miRNA-223 KO contributes to hypercholesterolemia and augmented accumulation of cholesterol in hepatocytes. However, Our data are only in consistent with the higher total cholesterol and LDL-C levels in livers from miRNA-223 conventional KO, ΔHep and KD_Hep_ mice, but failed to reproduce the hypercholesterolemia phenotype.

Our data support the strengthened cholesterol uptake and synthesis would be able to address the higher cholesterol contents in livers from miRNA-223 deficent mice but regulating mechanisms are not well identifcal with previously reports. On one hand, enhanced SR-BI expression in miRNA-223 deficent liver may promote the transportation of HDL-C from serum to hepatocytes as indicated by reduced HDL-C levels in serum. On the other hand, cholesterol synthesis would be enhanced in miRNA-223 ΔHep livers as evidenced by increased expression of active form of SREBP2, a key transcriptional factor regulating cholesterol synthesis. However, it seems a conflict raised between high cholesterol levels and the expression of SREBP2 and its directed genes, such as LDLR, in miRNA-223 deficent livers. Indeed, the balance of classical sterol feedback effects on SREBP2 activation fundamentally direct cholesterol synthesis or LDLR mediated uptake events in the liver cells; however, considering the “Time and Zone” concept for gene regulation, we thought our data from 5 weeks LD feeding would only reflect a specific period during Sterol-SREBP2-cholesterol synthesis feedback process within LD challenge. By unknown indirect regulation manner, miRNA-223 deficent would enhance hepatic SREBP2 expression and partially contribute to higher cholesterol content in liver, however as consequences of feedback from higher sterol levels, the activating effects of SREBP2 on targets gene expression were attenuated or less responsive as can be seen from only a few SREBP2 response genes (9 out of 15) would be mildly increased in ΔHepa livers in compare with cKO littermates, indicating the role SREBP2 is on the way of inactivation. As to “conflicting” LDLR expression in ΔHepa livers, we noticed that the elevated expression of proprotein convertase subtilisin kexin 9 (PCSK9), a crucial LDLR endosomal / lysosomal degradation protein was also identified, which might explain why LDLR protein levels were not consistent with increased SREBP2.

Since there was no significant hepatic injury occurred in ΔHep mice and their cKO littermates as indicated by comparable ALT and AST levels in serum, the next question was how the miRNA-223-deficient hepatocytes process the redundant cholesterol storage? We noticed lower serum levels of HDL-C and unaffected LDL-C in ΔHep mice when compared with their cKO littermates. However, the serum total cholesterol was comparable in both genotypes, suggesting additional source to replenish cholesterol difference. Indeed, in miRNA-223 ΔHep livers, we identified enhanced mRNA and protein expression of ABCA1, an important cholesterol efflux transporter to fulfill the circulating transportation of cholesterol from tissue or cells [27] which could promote cholesterol efflux to blood and facilitate hepatic cholesterol homeostasis. Vickers *et al*. reported that miRNA-223 indirectly up-regulates ABCA1 mRNA level and protein expression in Huh7 human hepatic cancer cell line via inhibiting the expression of transcriptional suppressor SP3, a direct target of miRNA-223 hence, SP1activation initiated the ABCA1 transcription[13]. Our data demonstrated that hepatocytes-specific loss of miRNA-223 enhanced ABCA1 mRNA and protein expression that is independent of SP1/SP3, as their expression was unaffected. Further investigation is required to resolve the controversial effects of miRNA-223 on ABCA1 expression in hepatocytes.

Our first novel finding is that miRNA-223 in hepatocytes plays a pivotal role in controlling biliary cholesterol secretion by direct targeting the canalicular cholesterol transporters i.e. ABCG5 and ABCG8. ABCG5/8 has been found to contribute ∼ 75% biliary cholesterol secretion, and its genetic mutation, or dysregulation would strongly disturbs cholesterol secretion and cholelithiasis progression. Our data solidly demonstrate that with LD challenge hepatocytes-specific miRNA-223 depletion significantly increases ABCG5 and ABCG8 expression and elevates cholesterol levels in both liver secreated bile and bladder stored bile. Meanwhile, we also detected increased SR-BI protein expression, which is believed to be an additional cholesterol transporter localized in canalicular membranes and performs cholesterol secretion role independent of ABCG5/8 in physical condtion[7], although its exact role in gallstone development remains unclear. Of the note, significant increased SR-BI expression evidenced in miRNA-223 hepatocytes specific KO livers, which is opposite with the previous reports in which reduced SR-BI expression appeared in miRNA-223 KO livers. The inconsistent observation perhaps derived from different strategys to generate miRNA-223 loss of functional mice or the specific experimental challenges.

Reversely, AAV-mediated hepatocytes-specific miRNA-223 OE in WT mice reduces expression of ABCG5,ABCG8 and decreases cholesterol content in the bile. In current study, we demonstrated a negative regulation manner between miRNA-223 and those cholesterol transporters in mouse hepatocyte. Although not noted in current available miRNA targets prediction databases, imperfect binding sequences of miRNA-223 were found in 3’UTR regions of *Abcg5* and *Abcg8* mRNA, suggesting a possible directly regulating manner. Therefore, hypothsis was experimental confirmed by classical 3’UTR reporter assays.

To our knowledge, conventional miRNA-223 KO mice were wildly used in various research fields, however, the tissue or cell specific miRNA-223 KO animal modle is emergently required to precisely verify the functional importance of miRNA-223 and molecular mechanism in interested cell type or tissue. Based on this view, we generated contional KO and consequently used conventional KO and cell specific KO of miRNA-223 in hepatocyte to clearly and logically demonstrate the crucial role of miRNA-223 in gallstone development and molecular mechanisms. We also suggested some retrospective studies should be considerd by using cell specific miRNA-223 KO mice, which whould betterly address the uncertain issues in previous studies by using conventional miRNA-223 KO animals.

Our second important finding is the novel role of miRNA-223 in cholesterol gallstone formation. Gallstone crystallization occurs in supersaturated bile where BA and PL cannot dissolve excessive cholesterol. Supersaturated bile could be attributed by 1) hyper-cholesterol secretion; 2) reduced biliary BA or PL secretion with unaffected cholesterol secretion; 3) increased cholesterol and reduced BA or PL secretion[3]. Our study suggested that accelerated gallstone formation in hepatic miRNA-223 KO/KD mice is strongly associated with supersaturated cholesterol status mainly attributed by an excess of cholesterol deposition in bile.

More interestingly, insufficient biliary PL levels were also determined in miRNA-223 deficient mice that would further decline cholesterol solubility, which also caused our interest. ABCG4 is known as a main PL transporter faciliateing hepatic PL secretion towards bile and its mRNA expression level was slightly increased in miRNA-223 deficent livers, it would be explained by a negative feedback way in which insufficient PL storage in bile requires more ABCG4 to increase biliary PL transportation. Similary, increased BA transporter ABCG11 in miRNA-223 deficent livers might attempt to balance the oversaturated cholesterol status in bile. It is a complexed gene regulating net work for PL and BA metabolism in hepatocytes. Although transcriptional factors, such as FXR, LXL and RXR, were important for PL and BA metabolism, their expression were not greatly affected in miRNA-223 deficent liver, suggesting other mechanisms and more detailed works needed in the future studies to determine the potent role of miRNA-223 in regulating hepatic PL and BA homeostasis.

Besides, recent research also indicates that gallbladder motility insufficiencyisgreatly affected by the loss of interstitial Cajal-like cells and telocytesthatfurther contributes to the cholelithiasis [28, 29]. Apparently, whether hepatic miRNA-223-affected bile saturation might have association with gallbladder motilityd ysfunction (specially Cajal-like cells and telocytesloss) is an interesting question and worthwhile to explore in the future study.

Lastly, this study provides the evidence for the miRNA-223 to be an effective therapeutic target in cholesterol gallstone disease. To evaluate the therapeutic efficacy by manipulating miRNA-223 expression, we conducted AAV-mediated miRNA-223 overexpression in WT mouse liver. The results demonstrated that elevating miRNA-223 levels in hepatocytes efficiently decreased bile cholesterol content and reduced gallstone formation, supporting a novel therapeutic strategy for cholesterol gallstone disease by targeting miRNA-223.

In summary, the present study uncovers an important role of miRNA-223 in regulating hepatobiliary cholesterol secretion and cholesterol gallstone formation by targeting key cholesterol transporters ABCG5 and ABCG8 in cholesterol biliary secretion pathway. Proof-of-concept study using AAV-mediated miRNA-223 OE in WT mouse liver shows promising therapeutic efficacy in treating LD-induced cholesterol gallstone. Hence, miRNA-223 has been identified as a novel potential therapeutic target in cholesterol gallstone disease.

## Abbreviations

miRNA-223: microRNA-223;
LD: lithogenic diet;
ABCG: ATP-binding cassette sub-family G;
T-Chol.: total cholesterol;
BA: bile salt;
PL: phospholipids;
TG: triglyceride;
PHC: primary hepatocytes;
BM: bone marrow.

## Acknowledgments

This study was supported by the National Natural Science Foundation of China (81570236, 81870360, and 81702626), Liaoning Pandeng Scholar Program (To Prof. Tonghui Ma), Liaoning Provincial Program for Top Discipline of Basic Medical Sciences.

## Author contributions

Research design: Lei Shi, Tonghui Ma

Conducting experiments:Feng Zhao, Shiyu MA, Wei Shen, Xiaolin Cui, Lina Xing, Zihe Peng,Xiang Li.

Yanghao Li, Gang Liu

Data acquisition: Zhao Feng, Shiyu Ma, lingling Jin

Data analysis: Shiyu Ma, Feng Zhao

Writing the manuscript: Lei Shi, Tonguhi Ma, Feng Zhao, Wei Shen

## Supplementary Figures

**Supplementary Figure 1.**
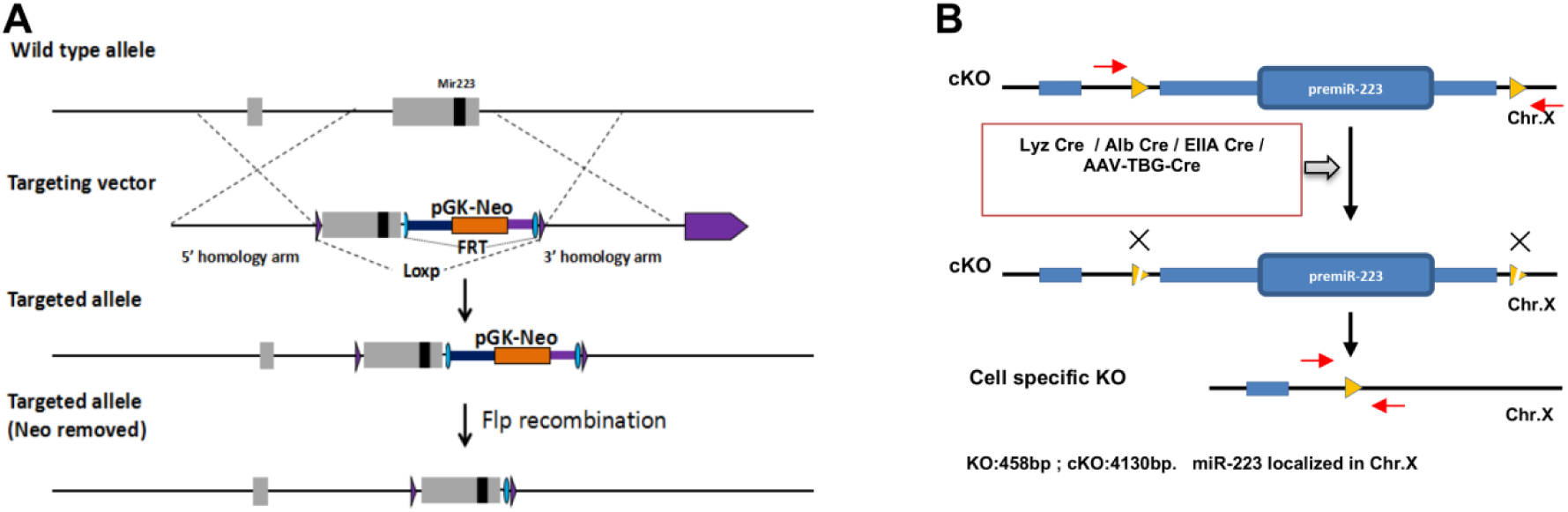
Strategys of generating MiRNA-223 conditional knockout and cell specific knockout / knockdown mice stains. A. miRNA-223(ENSMUST00000102112) was localized in the second Exon of ncRNA F630028O10Rik, two Flox sequences were designed in the Intron 1 and down stream of Exon 2. By homologue recombinant targeting principle in embryonic stem cells, targeting vector were prepared containing 9.616 kb 5’ homologue arm, 3.566 kb Flox region, PGK-Neo-polyA region flanked with FRTs, 4.919 kb 3’ homologue arm, and MC1-TK-poly A negative selection region. The positive EC clones were obtained by G418 and Ganc selection and genomic DNA sequencing verification, and chimeric offsprings were obtained after implantation of C57BL/6J blastaea injected with recombinant ES cells into surrogate mice. Neo cassette was removed by breeding chimeric mice with Flp^Tg^ mice. All founder mice were verified by genomic DNA sequencing. B Scheme showing the stragte for generating tissue or cell specific miRNA-223 KO mice by crossbreeding miRNA-223 cKO mice with indicated Cre mice or virus infection.

**Supplementary Figure2.**
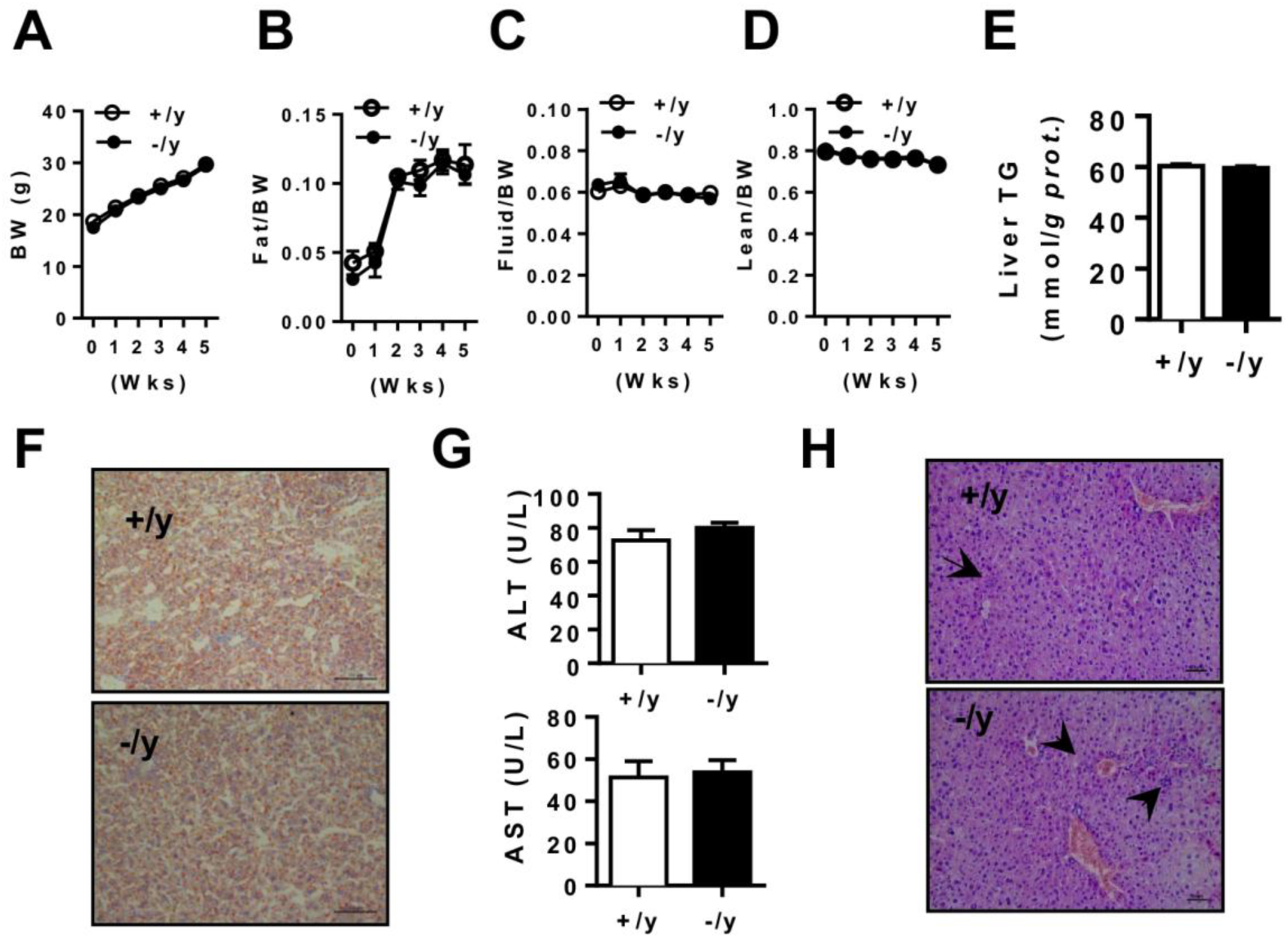
During 5 weeks LD treatment, (A) body weight changes, (B) Fat contents, (C) Fluid contents and (D) Lean contents were evaluated in the indicated time for in +/y and -/y mice (n=5 mice each group). After 5 weeks LD feeding, +/y and -/y mice liver tissues (8 mice each group) were collected for (E) TG contents; (G) ALT and AST activities in serum were determined, (H) H&E staining for liver sections and arrow indicated the infiltrating inflamatory cells.

**Supplementary Figure3.**
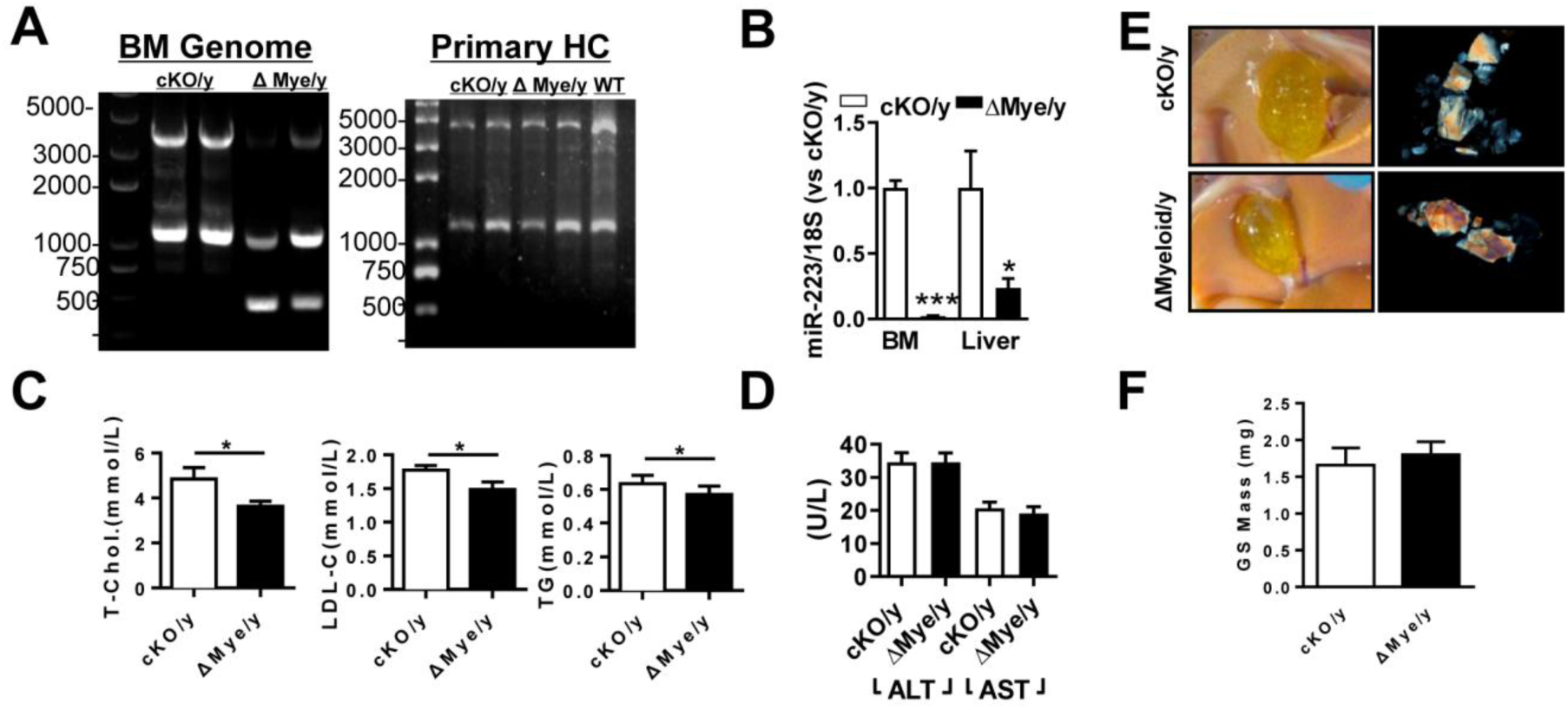
(A) PCR showing the genomic DNA precision for miRNA-223 loci in Bone marrow (BM) and primary hepatocytes (HC), and (B) RT-qPCR revealed miRNA-223 epxression levels in BM and HC from miRNA-223 KO mice (Δmyeloid/y) and cKO/y mice (cKO.+/y:4130bp, KO:458bp, n=4 mice each group); With 5 weeks LD treatment, the levels of (C) Total-Cholesterol (T-Chol.), LDL-C and TG in livers and (D) Serum ALT and AST acitivities were determined (n=10 each group); (E&F) gallstone phenotype and gallstone mass were further determined (n=4 each group). \**p*<0.05; \*\*\**p*<0.001 versus cKO/y.

**Supplementary Figure 4.**
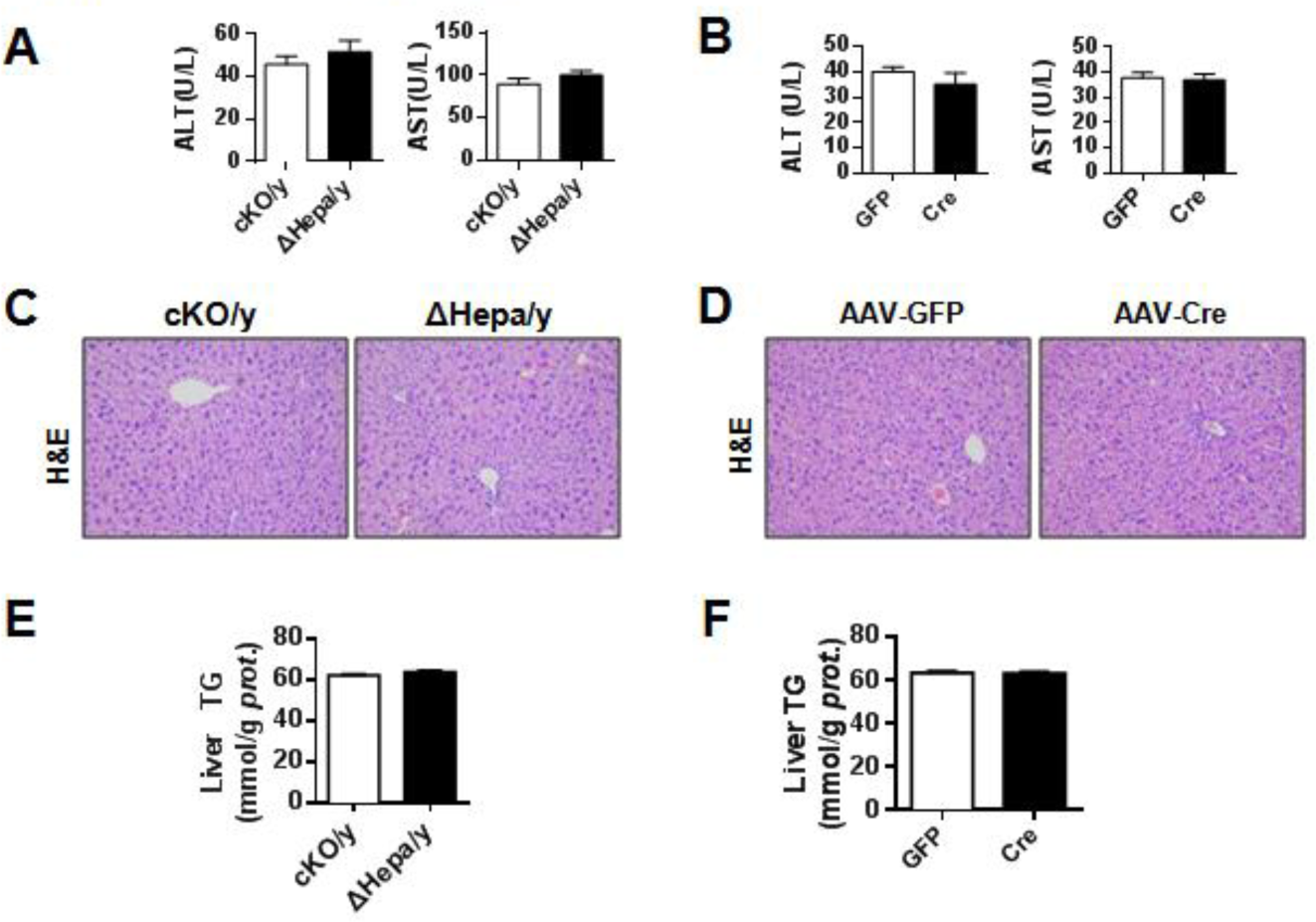
The effects of hepatocyte specific miRNA-223 KO or KD on liver injury and TG content.ΔHepa/y and cKO/y mice (n=10 mice each group) or AAV-TBG-Cre and AAV-GFP treated mice (n=6-8 mice each group) were fed with LD for 5 weeks, (A&B) serum were used to determined ALT and AST activities, liver tissues were analyzed for (C&D) H&E staining and (E&F) TG contents.

**Supplementary Figure 5.**
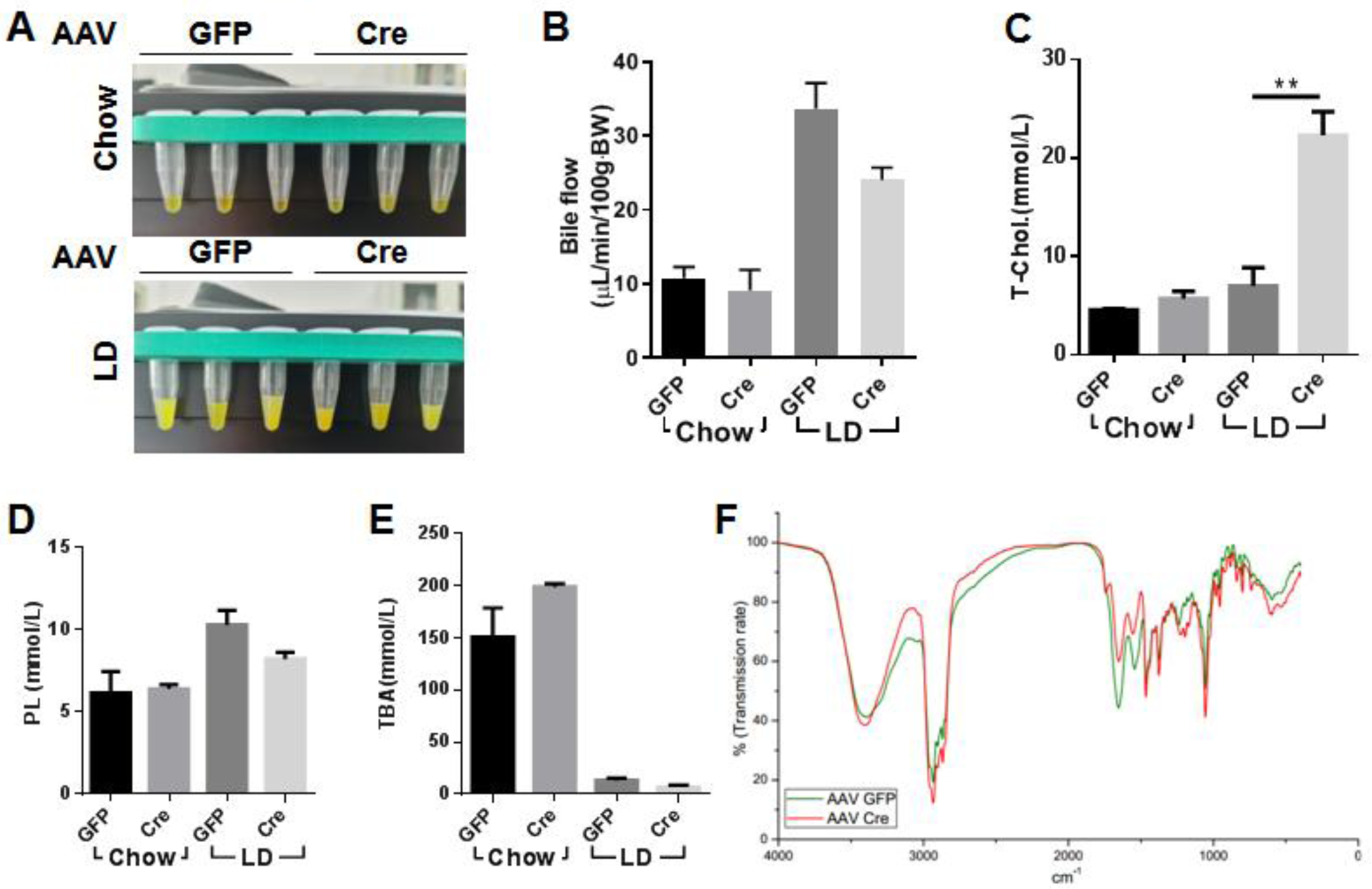
The effects of miRNA-223 knockdown on mice bile secretion and lipid contents in bile and gallstone. Hepatocyte specific miRNA-223 knockdown were conducted via AAV8-TGB-Cre or AAV8-CMV-Cre (control) *i*.*v*. injection for miRNA-223 cKO/y mice and three weeks later, those mice were fed with chow or LD for additional three weeks. Bile flow rate was determined via catheterization with PE-10 tube within 30 min. (A) representative image showing the liver secreted bile volume; (B) Bar graph showing bile flow rate Bile lipids content of (C) T-Cholesterol, (D) PL and (E) TBA were separately determined from liver secreted bile. Data are summarized from 3-4 mice each group and \*\**p*<0.01 versus GFP. (F) The affects of miR-223 KD on cholesterol contents in mice gallstone. Gallstones were collected and mixed with KBr (1:100) and further prepared for Fourier Infra-red Spectrograph analysis. The cholesterol specific regions are at 2960.244, 2935.173, 2902.383, 2867.676 cm^-1^. Gallstones from 3 mice each group were mixed and subjected to Fourier Infra-red Spectrograph analysis.

**Supplementary Figure 6.**
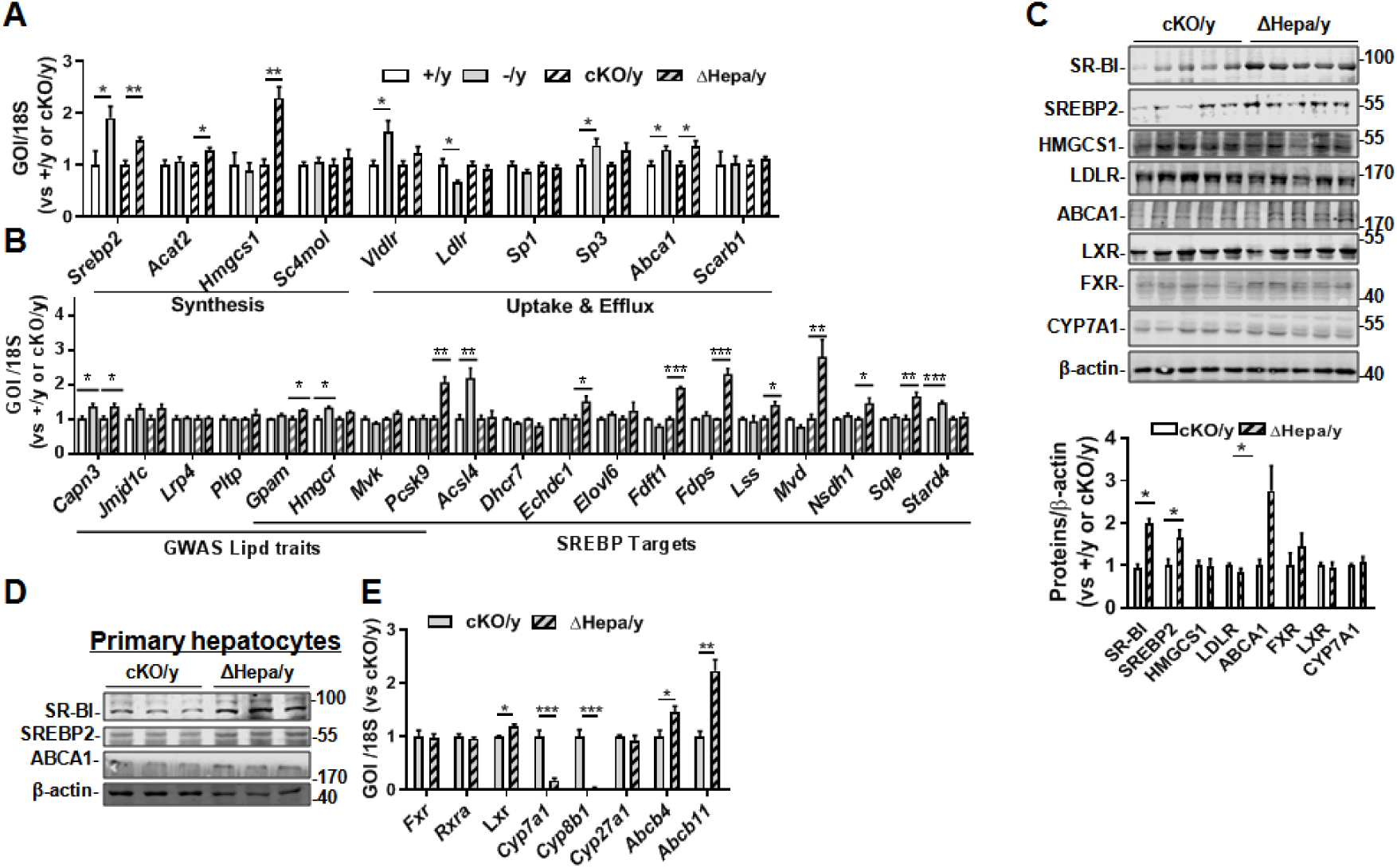
The gene expression was determined by RT-qPCR or western blotting in livers samples from -/y and +/y or ΔHepa/y and cKO/y mice with LD challenge for 5 weeks. (A) RT-qPCR assessed the selected genes expression concerning hepatic cholesterol synthesis, uptake and efflux as well as biliary secretion (n=5-6 mice per group). (B) RT-qPCR evaluated the mRNA expression levels as indicated by GWAS lipid traits and SREBP2 targets (n=5-6 mice per group). (C) Liver protein levels for SR-BI (∼75 kDa), SREBP2 (∼55 kDa), HMGCS(∼52 kDa), LDLR(∼160 kDa), ABCA1(∼250 kDa), LXR(∼45 kDa), FXR(∼59 kDa) and CYP7A1(∼50 kDa) as well as (D) primary hepatocytes expressing SREBP2, SR-BI and ABCA1were determined by Western blotting (n=3-5 mice per group). (E) RT-qPCR exams the mRNA expression for indicated genes from livers of cko/y and Δhepa mice (n=5-6 mice per group). \**p*<0.05, \*\**p*<0.01 or \*\*\**p*<0.001 versus +/y or cKO/y.

**Supplementary Figure 7.**
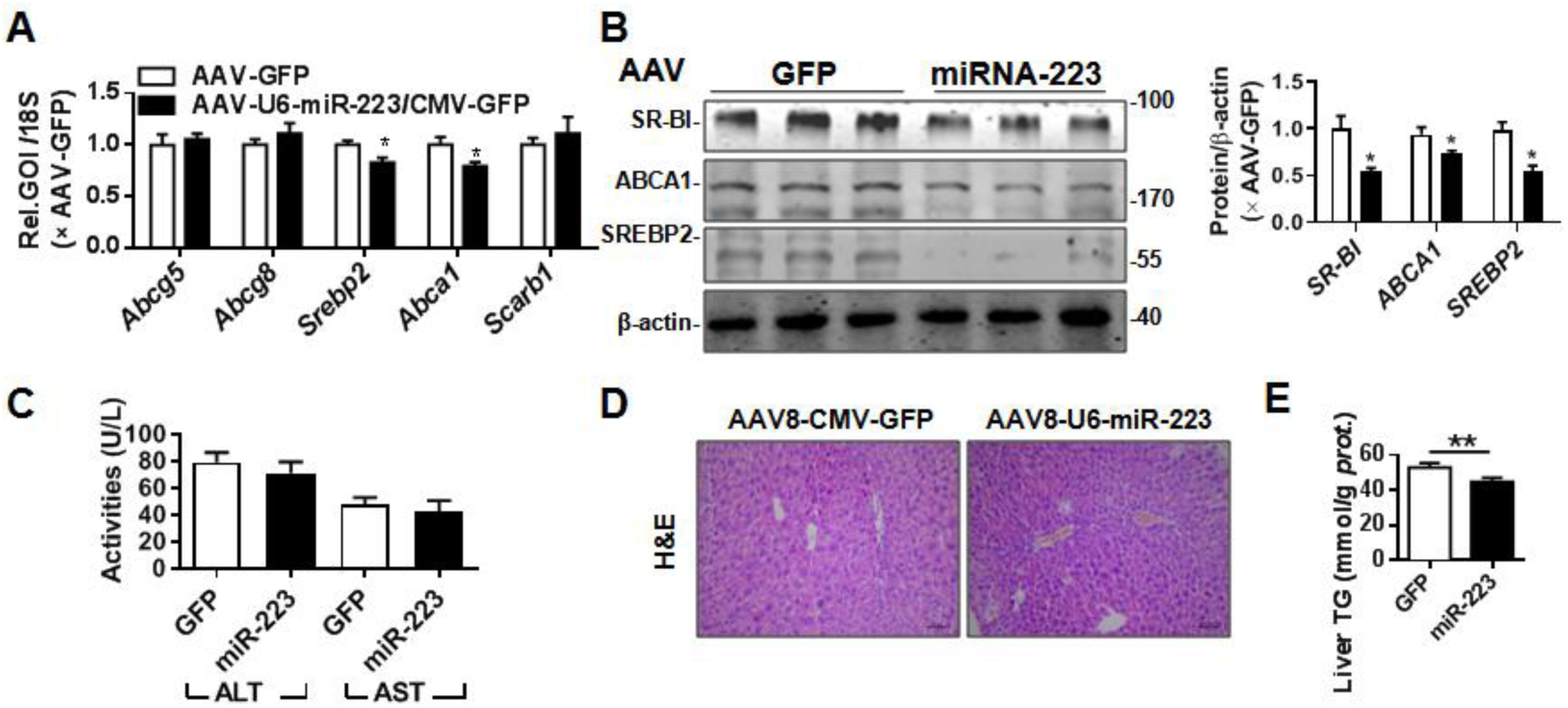
WT mice were pretreated with LD for 3 weeks followed by one time injection with AAV8-U6-miRNA-223/CMV-GFP or AAV8-CMV-GFP (10^11^ virus genome) and continued LD feeding for additional 5 weeks, (A) mRNA expression of *Abcg5, Abcg8, Srebp2, Abca1* and *Scarb1* were determined by RT-qPCR *(n=5 mice per group)*; (B) protein levels of SR-BI, SREBP2 and ABCA1 were detected by western blotting (n=3 mice per group); (C) serum ALT and AST activites (n=9 mice per group); (D) H&E staining for liver sections and (E) liver TG levels (n=9 mice per group). \**p*<0.05, \*\**P*<0.01 versus GFP.

## Supplementary Lists

**Supplementary list 1.**
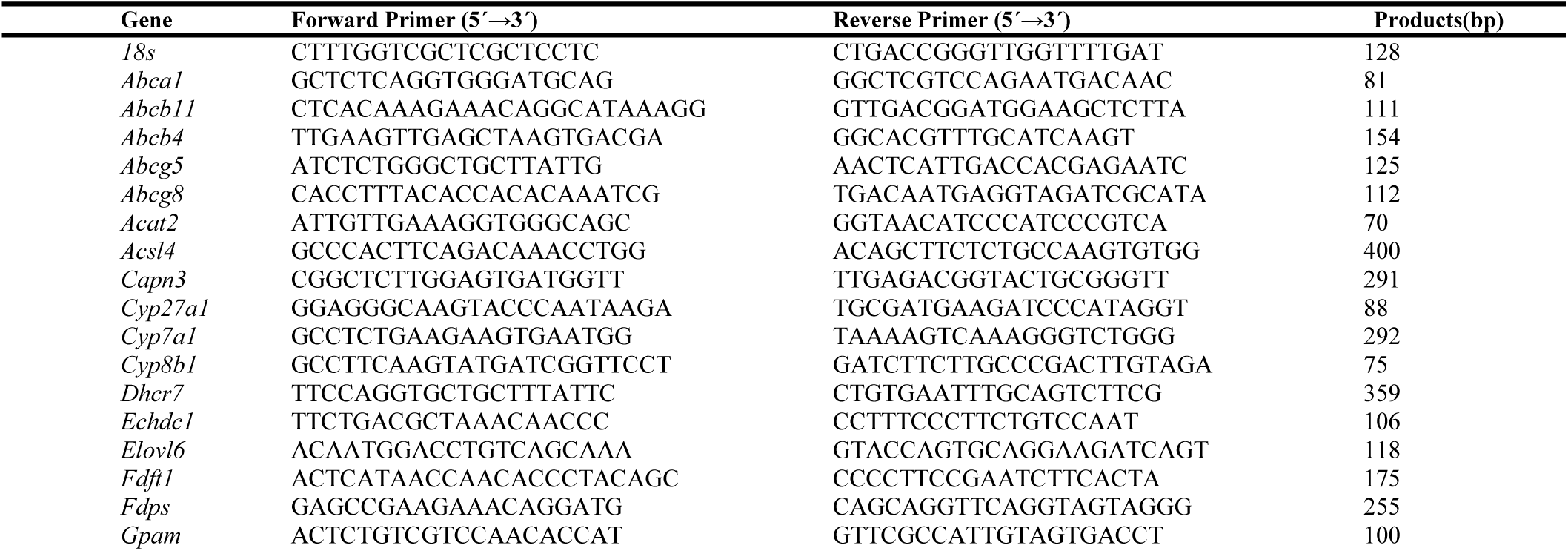
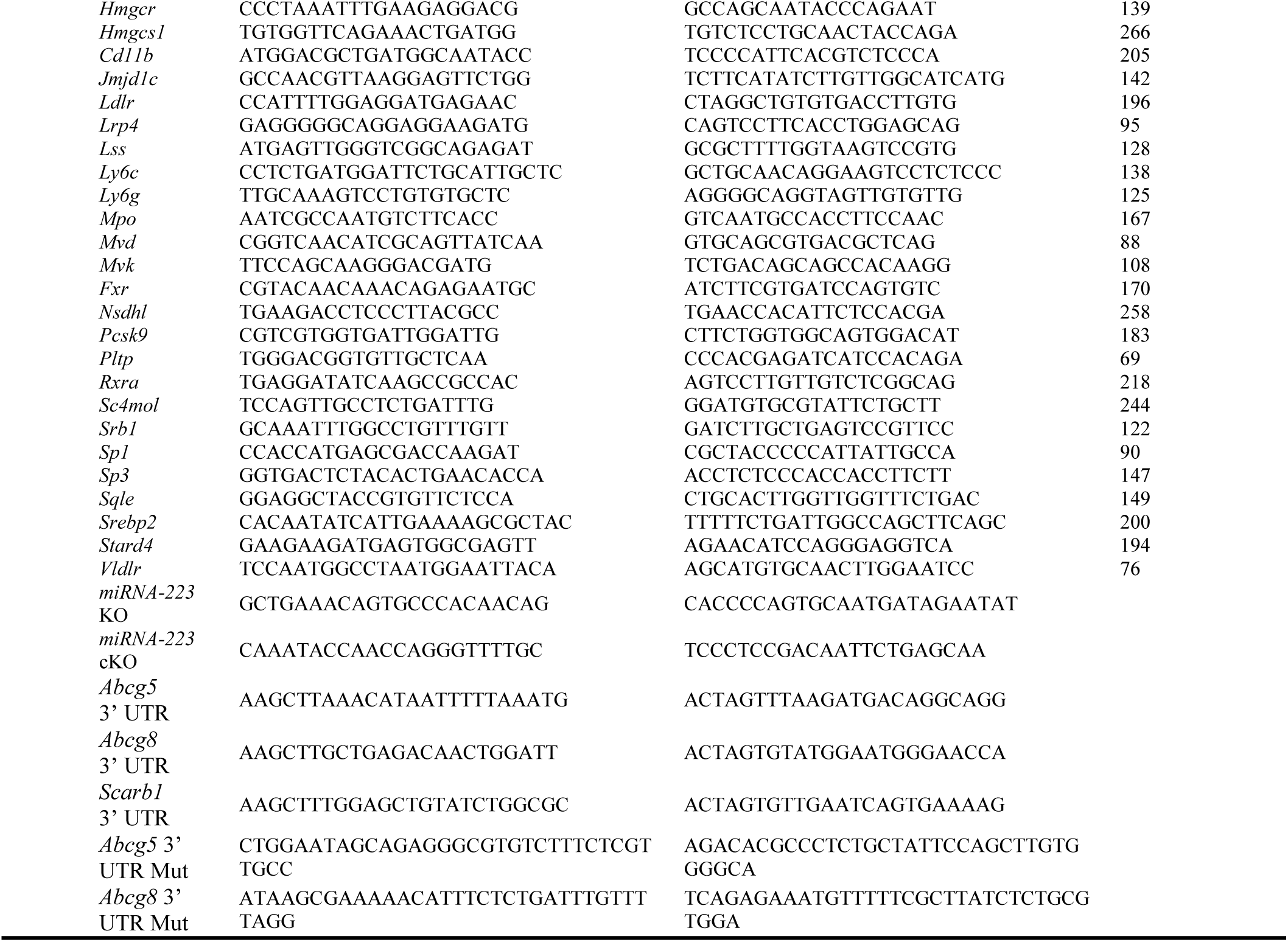
Forward and Reverse Primers Used for RT-qPCR, Genotyping and Plasmid construction

**Supplementary list 2.**
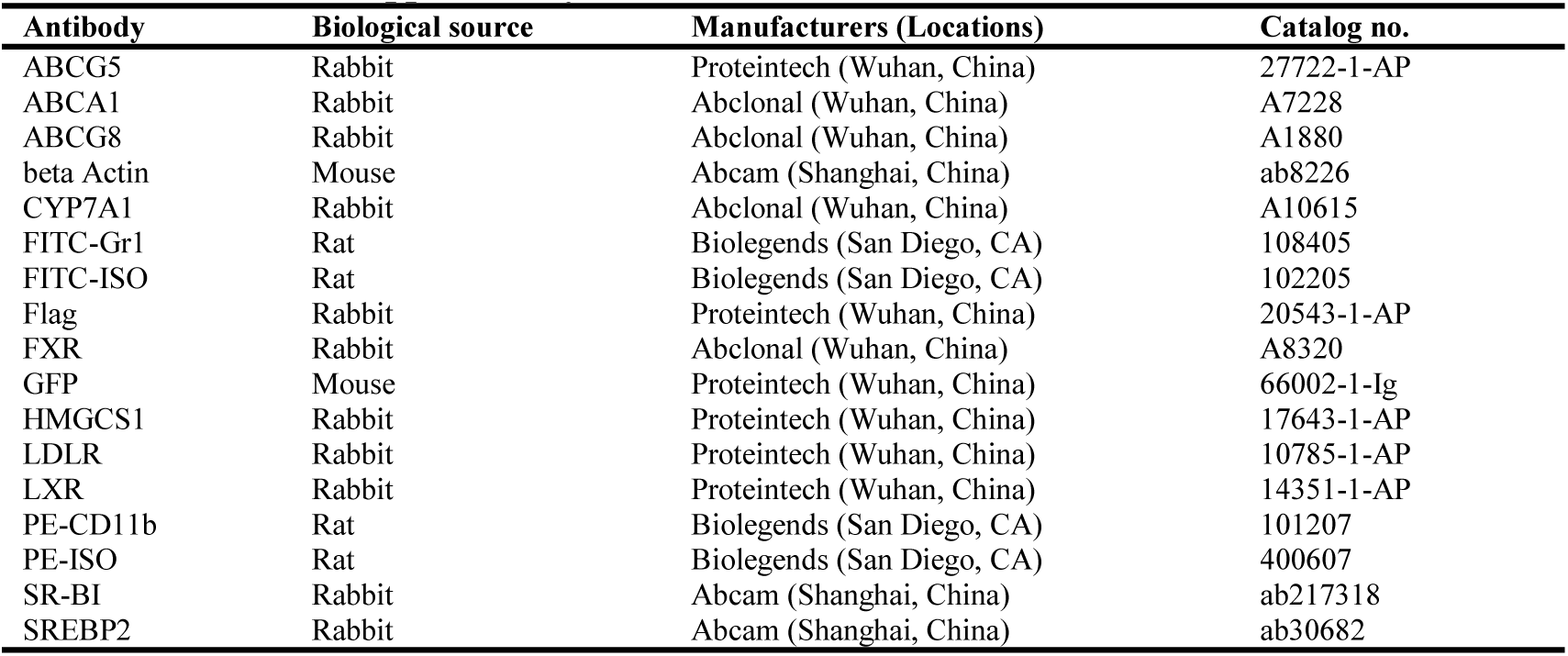
Antibodies Used for Western Blot and FACS

